# Designing host-associated microbiomes using the consumer/resource model

**DOI:** 10.1101/2023.04.28.538625

**Authors:** Germán Plata, Karthik Srinivasan, Madan Krishnamurthy, Lukas Herron, Purushottam Dixit

**Author notes:** Contributed equally (Germán Plata, Karthik Srinivasan).

## Abstract

A key step towards rational microbiome engineering is *in silico* sampling of realistic microbial communities that correspond to desired host phenotypes, and vice versa. This remains challenging due to a lack of generative models that simultaneously capture compositions of host-associated microbiomes and host phenotypes. To that end, we present a generative model based on the mechanistic consumer/resource (C/R) framework. In the model, variation in microbial ecosystem composition arises due to differences in the availability of *effective resources* (inferred latent variables) while species’ resource preferences remain conserved. The same latent variables are used to model phenotypic states of hosts. *In silico* microbiomes generated by our model accurately reproduce universal and dataset-specific statistics of bacterial communities. The model allows us to address three salient questions in host-associated microbial ecologies: (1) which host phenotypes maximally constrain the composition of the host-associated microbiomes? (2) how context-specific are phenotype/microbiome associations, and (3) what are plausible microbiome compositions that correspond to desired host phenotypes? Our approach aids the analysis and design of microbial communities associated with host phenotypes of interest.

## Introduction

Host-associated microbiomes play important roles in host health and disease [1] and, for livestock species, feed efficiency and environmental footprint [2]. Unfortunately, however, rational design of microbiomes to achieve desired host phenotypic states remains challenging due to extensive variability [3-5], high dimensionality, and dynamic nature of microbiomes [4, 6, 7], co-evolution of hosts and microbiome [8, 9], and numerous microbe ↔ microbe [10] and host ↔ microbe interactions [8, 9, 11].

A key step in rational design of high dimensional biological systems is generative modeling. Generative models are machine learning models that probabilistically generate *in silico* data and reproduce key features of experimental observations [12]. These models have been successfully used to model protein sequences [13-15], genomes [16], presence/absence patterns in microbial ecosystems [17], and single cell transcription profiles [18]. Generative models have been used for data augmentation and to identify functional covariation in the data (e.g. residue-residue contacts in proteins [19]). Generative models can assign probabilities to unseen observations and therefore can be used, for example, to identify disease effect of genetic variants [20]. Importantly, generative models have been used to design biological systems with desired properties, for example, proteins with higher enzymatic activity [13]. Generative models can be extremely useful for microbiome engineering, for example, in identifying feasible microbiome community structures that correspond to desired community functions or host phenotypes or in identifying specific host phenotypes, environmental parameters, or microbial taxa that constrain community composition.

Off-the-shelf generative models based on generative adversarial neural networks [21-24], maximum entropy methods [25], and mixture Dirichlet distributions [26] have been used to study the composition of microbiomes. However, many of these models tend to have a very large number of parameters and run the risk of overparameterization when modeling microbiomes. This is because unlike other biological data (protein sequences, RNAseq etc.), microbiome studies usually comprise significantly smaller sample sizes (*N* ∼ 100) and exhibit extensive variation. Additionally, these generative models provide little insight into the environmental [27, 28] and host factors [29] that drive the diversity in microbiomes. Notably, state-of-the-art generative models in microbiome studies have not modeled the simultaneous variation in corresponding host phenotypes.

An alternative to data-driven generative modeling is mechanistic generative modeling. Here, variation in microbiome composition is explained using variation in interpretable and mechanistic parameters of the model. The consumer/resource (C/R) framework is one of the most popular mechanistic models to understand variation in microbiome composition [28, 30-36]. In C/R models, variation in microbiomes arises due to variation in resource availability while the preferences of microbial species (consumers) towards these resources remain conserved across similar ecosystems. With an appropriate choice of parameters, ecosystem compositions generated using the C/R model reproduce many universal statistical features of real and synthetic microbiomes [28, 32, 36, 37] such as distributions of diversity metrics [28] and scaling relationships [27]. Unfortunately, fitting the C/R model to data is difficult as it requires knowledge of high-resolution temporal dynamics of species abundances, or the concentrations of resources consumed by microbial species through time [38] or species-resource preferences [39]. Consequently, training these models on specific microbiome datasets is difficult even for controlled *in vitro* communities [36]. Therefore, studies using the C/R model cannot model covariation patterns observed in specific host-microbiome systems, instead randomly drawn parameters have been used to explain statistical features of the data [27, 28]. Finally, the C/R framework is not able to model variation in host phenotypes, except for the abundances of metabolites directly consumed/excreted by the microbiome.

Here, we develop a generative model for host-associated microbiomes using the mechanistic C/R framework. To that end, we identify a latent space induced from the C/R model that encodes the temporal histories of *effective resources* affecting microbial growth. Variation along this latent space results in variation in microbiome compositions. Using data from three host-associated microbiomes, we show that probabilistic modeling of this latent space reproduces both universal statistical features as well as system-specific covariations between bacterial taxa. Using the model and the bovine rumen microbiome as a test case, we investigate three salient questions in microbiome design and engineering: (1) which host phenotypes maximally constrain the composition of the host-associated microbiome? (2) how context-specific are phenotype/microbiome associations, and (3) what are plausible microbiome compositions that correspond to desired host phenotypes?

To the best of our knowledge, this is the first generative model that can simultaneously model microbiome compositions and the environmental factors, for example, phenotypic states of the host. Therefore, we believe that our generative model will be an important tool for the design of host associated microbiomes.

## Results

### The C/R framework identifies a latent space to build a machine learning model

We use the C/R framework to identify a latent space to model microbiome compositions (SI Section 1, Figure 1). In the C/R model, the change in abundance of an organism (growth rate) depends on the availability of resources *r*_*k*_ and species preferences towards those resources *θ*_*ko*_. The resources themselves are depleted according to their consumption by all organisms and net flow. As discussed above, fitting the C/R model to data requires not only the knowledge of microbial abundances but also the identities and abundances of those resources, either across samples or over time. Typically, these measurements are not available and, consequently, the C/R model is only used for its qualitative insights. Here, we show how to convert the C/R model to a latent variable model to describe species abundances. To do so, we integrate the equation describing the growth rate of a species in the C/R model and obtain an expression for the species abundance *n*_*o*_(*t*) at time *t*:

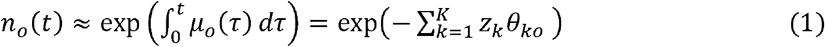

where *μ*_*o*_(*τ*) is the time-dependent growth rate. We recognize:

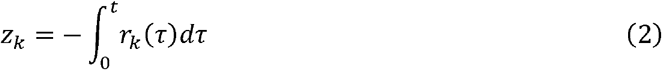

as latent variables that represent the complex temporal dynamics of resources. We note that the latent variables thus defined are emergent variables; they correspond neither to the initial concentration nor to the steady state concentrations of the resources. Instead, they represent the temporal history of resource fluctuations which in turn may depend on flow rates, resource sharing, and cross-feeding [31, 37].

**Figure 1.**
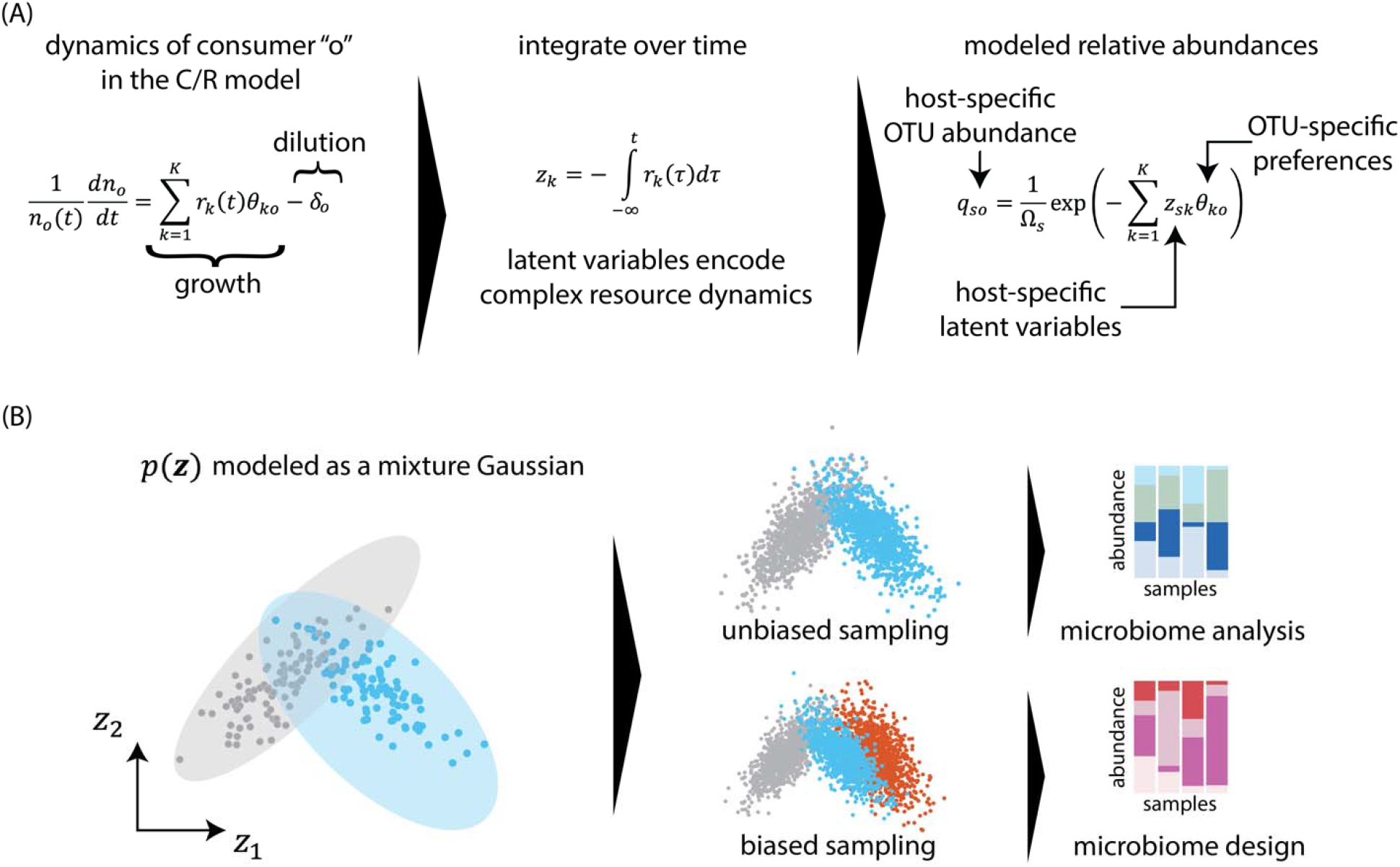
A generative model for host-associated microbiomes using the C/R framework. The C/R model defines changes in the abundance *n*_*o*_(*t*) of consumers “o” as a function of resource abundances *r*_*k*_(*t*) and consumer preferences *θ*_*ko*_ towards resources. Integrating the C/R dynamics over time shows that species abundances are a function of latent variables *z*_*k*_(*t*) that represent the sample-specific time history of resources, and universal species-specific preferences towards those latent variables. (B) To generate realistic microbiomes, sample embeddings in the latent space are modeled using a mixture Gaussian distribution. The model allows data augmentation via unbiased sampling in the latent space. Biased sampling from the model allows us to generate microbiome samples that correspond to desired host phenotypes.

Recognizing these latent variables allows us to circumvent the numerical difficulties in fitting the C/R model to microbiome data [39]. Observe that the relative abundances of organisms *q*_*o*_ are simple exponential functions of the latent variables:

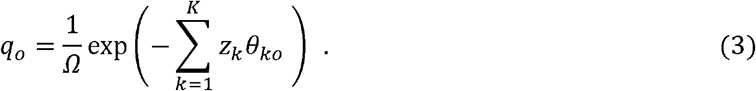

Unlike the C/R model, Eq. 3 can be easily fit to cross sectional microbiome data [40]. To that end, consider that the relative abundances (relative counts) *x*_*so*_ of *O* OTUs (operational taxonomic units) are measured in *S* hosts. We assume that the resource preferences of OTUs are universal across hosts while resource dynamics are host dependent. That is, latent variables *z*_*sk*_ are host-specific but species-independent and *θ*_*ko*_ are species-specific but host-independent. For fitting Eq. 3 with a fixed *K*, we minimize the cross-entropy *C* [36], which is equivalent to maximizing a probabilistic model for species abundances that follows a multinomial distribution (Eq. 4).

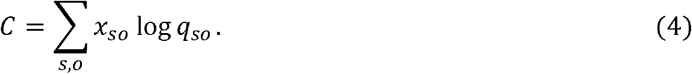

As we show in SI Section 1, this minimization is a nonlinear low rank matrix factorization wherein and *θ* can be learnt using gradient descent.

Unlike other machine learning models, our model has clear mechanistic interpretation: microbial couplings to specific latent variables remain constant across different ecosystems, but the temporal histories of these latent variables vary from ecosystem-to-ecosystem, leading to observed variation in community compositions.

### Host-associated microbiomes are surprisingly low-dimensional

How accurate is Eq. 5 in modeling compositions of microbiomes? We studied microbiomes associated with three host species, the rumen of Holstein cows [41], the chicken cecum [42], and the human gut [43]. The bovine and the chicken microbiomes were characterized at the level of operational taxonomic units (OTUs) while the human microbiomes were characterized at the genus level. Based on our previous analysis on technical noise associated with sequencing of microbial samples, we only included taxa whose average abundance was higher than 0.1% [5]. In the three datasets, there were 790 samples and 157 OTUs (cows), 778 samples and 156 OTUs (chicken), and 127 samples and 95 genera (humans) respectively (**SI Section 2**).

The number of resources consumed by species in these ecosystems can potentially be very large (*K* ∼ 10^3^ − 10^4^) [36]. To avoid overparameterization in our model, we seek a small number of *effective resources* (latent variables) [44] that capture the observed compositional variation of microbiomes. These latent variables do not correspond to specific nutrients present at a specific time. Instead, they represent a combination of factors that both positively and negatively affect the growth of microbial species aggregated over time. To determine the effective dimension of the latent space, we fit Eq. 5 where *K* is chosen as the smallest of the number of hosts *S* and the number of species *O*. We determined the effective dimension by examining the singular values (SVs) of *Z* × *Θ* (insets of Figure 2ABC). Singular values quantify the variance explained by low rank approximations to matrices and therefore can identify coarse-grained latent variables that can accurately approximate the microbiome. SVs for all three ecosystems showed two regimes. When arranged from largest to the smallest, SVs first decreased rapidly, followed by a gradual power law-like decrease. This suggested that a small number of latent variables captured the essential variation in microbial composition. To confirm this, we fit microbiome compositions using different values of *K*. Indeed, as seen in Figure 2DEF, the symmetric Kullback-Leibler divergence and the Bray-Curtis dissimilarity between data and model fits decreased approximately logarithmically with increasing dimension of the latent space and at a significantly slower rate for larger *K*s.

**Figure 2.**
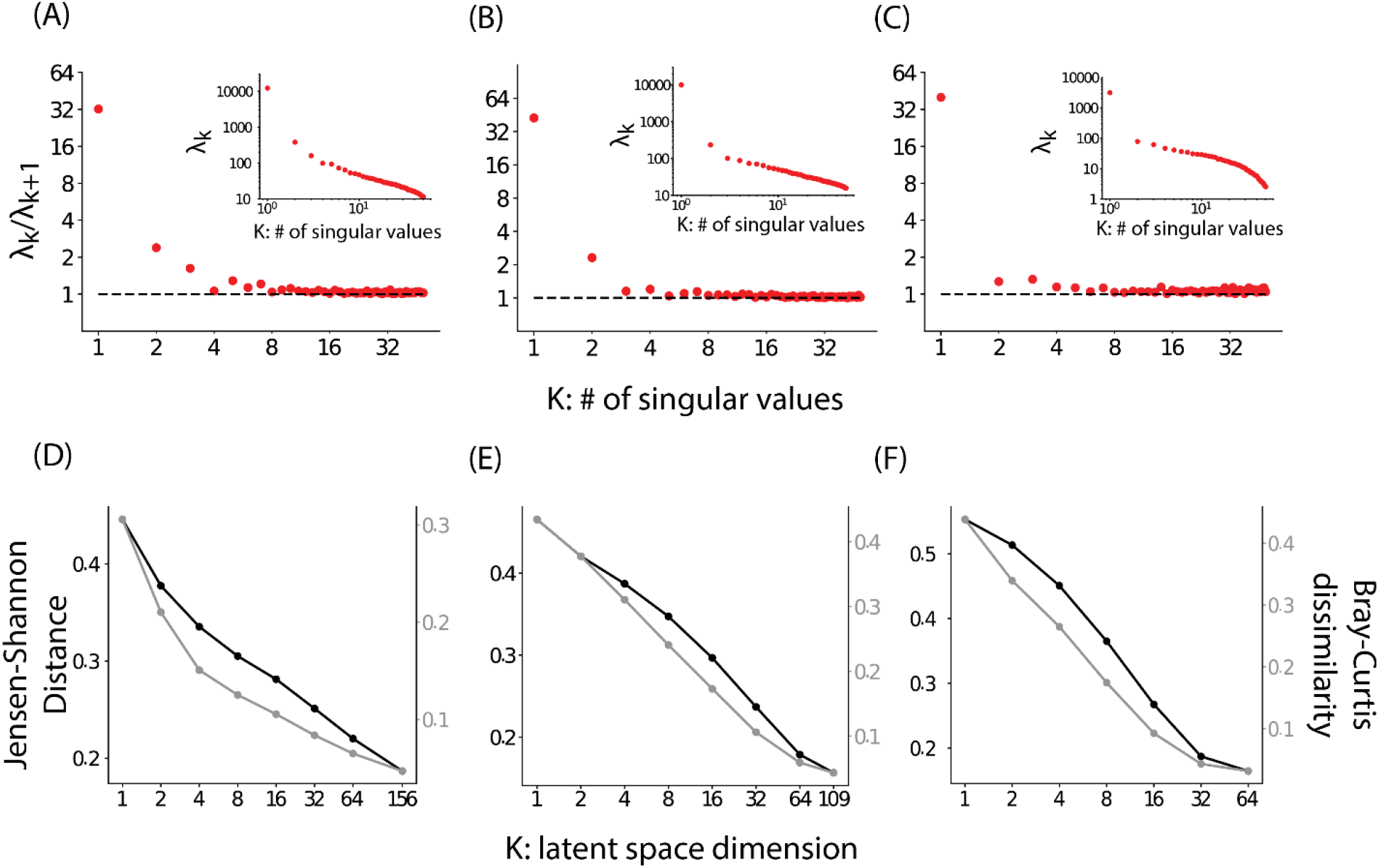
Low dimensionality of host-associated microbiomes. Panels (A), (B), and (C), the ratio of successive singular values 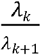 of the *Z* × *Θ* matrix for bovine, chicken, and human microbiomes respectively. Only the first 50 SVs are shown as later, less important SVs may be dominated by noise. The insets show the singular values arranged in a decreasing order. Panels (D), (E), and (F) show the average symmetric Jensen-Shannon distance(black) and the Bray-Curtis dissimilarity (gray) between observed community composition and the corresponding model fit as a function of the dimension of the latent space. The average is taken over all samples.

These results show that a low dimensional latent space constructed using the C/R model captures compositional variation in microbiomes in three different host species. We note that this dimensionality reduction approach is conceptually different from principal coordinate analysis (PCoA), a standard tool in microbiome analysis. In PCoA, one approximates dissimilarity between compositions of pairs of ecosystems using dimensionality reduction obtained via principal component analysis performed directly on the pairwise distance matrix. The variance explained by PCoA is therefore the variance explained in predicting pairwise distances using a lower dimensional model. In contrast, our approach directly models (approximates) compositions of individual ecosystems using a lower dimensional model. Notably, the accuracy of our approach quantified in terms of the variance explained in reconstructing community composition as well as reconstructing pairwise Bray-Curtis dissimilarity shows that a small number of components is sufficient to capture both community composition as well as *β* − diversity (SI Figure 1).

This low dimensionality may arise due to covariation in species that results from specific structure in consumer/resource matrices [33, 35], due to covariation in resource flow rates [33], due to universal factors such as pH [45], or due to coarse-grained structure/function relationships between the microbiome and its environment [34] that affect broad classes of species in a similar manner. Notably, as we show below, this low dimensionality was not due to low compositional diversity. Indeed, species abundances were widely distributed with an abundance distribution (Figure 3C) that had an extended power law scaling between abundances of 0.1% to 10%. In this work, we exploit this low dimensionality for generative modeling. We leave it for future studies to explore its mechanistic underpinnings.

**Figure 3.**
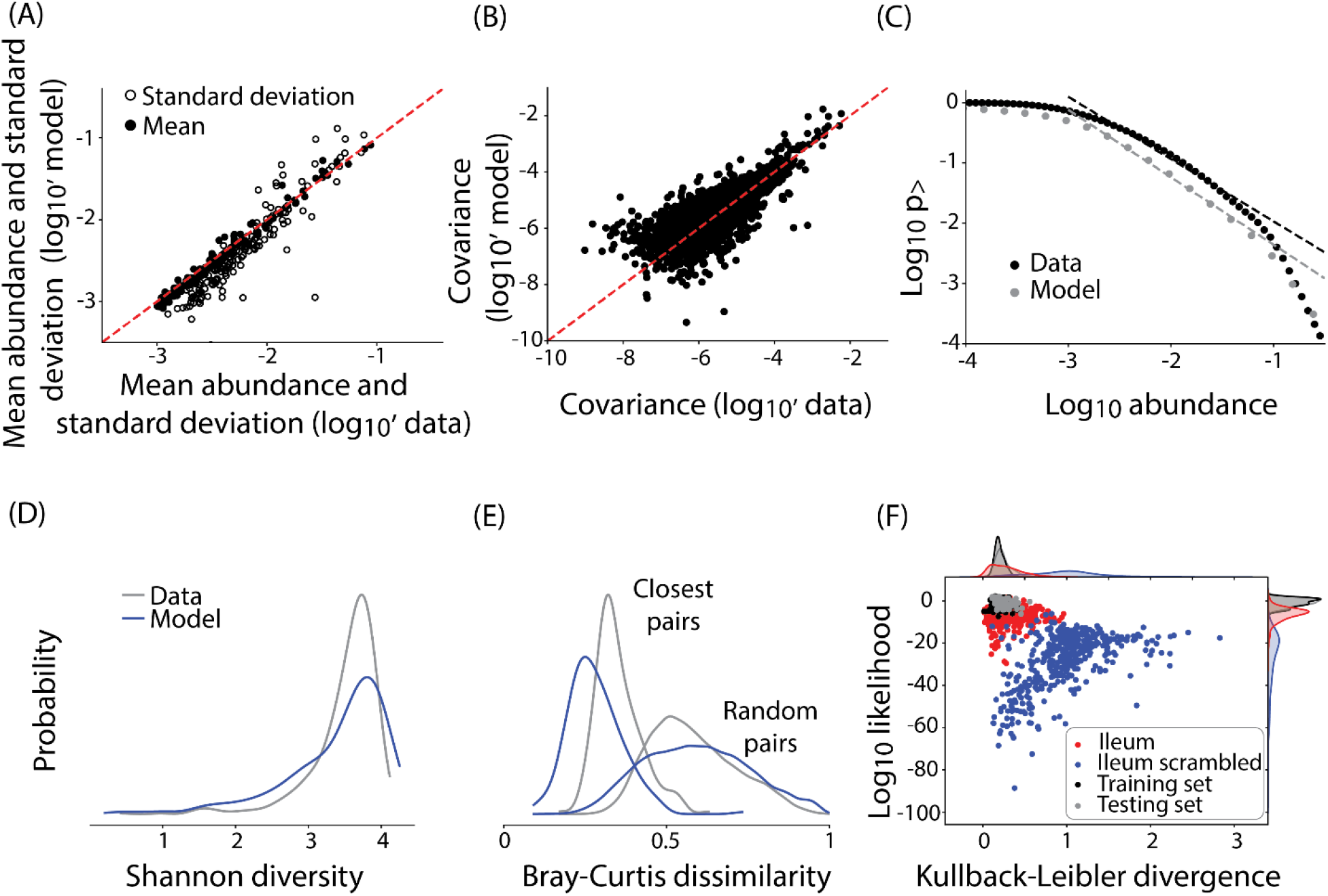
Latent space-based model generates accurate *in silico* communities. (A) A comparison between mean OTU abundances (filled circles) and standard deviations (open circles) and the corresponding model predictions (y-axis). The dashed red line represents *x* = *y*. (B) A comparison between OTU-OTU covariance ⟨*δx*_*i*_*δx*_*j*_ ⟩as computed in the data and as predicted by the model. Absolute values are shown. (C) The inverse cumulative distribution representing the probability of observing an OTU abundance greater than a given abundance (x-axis) as computed from the data (black) and the corresponding model prediction (gray). The dashed lines show the best fit power law between relative abundances of 10^−3^ and 10^−1^. The slope of the power law is −1.03 in the data and −1.12 in the model. (D) The distribution of Shannon diversity of community compositions as observed in the data (gray) and computed from the model generated communities (blue). (E) The distribution of Bray-Curtis dissimilarity between random pairs of community as well as closest pairs of communities in the data (gray) and the model (blue). Distributions are smoothed using Gaussian kernel density estimation. (F) a scatter plot of Kullback-Leibler divergences between community composition in the data and the corresponding model fit (x-axis) and the log-likelihood of the embedded latent variables (y-axis). Data are shown for the training data from chicken cecum (black), testing data from chicken cecum (gray), testing data from chicken ileum (red), and testing data from chicken ileum embedded using a scrambled preference matrix (blue).

Finally, we confirmed that the trends in community reconstruction accuracy did not depend on the abundance cutoff used as the OTU inclusion criteria (SI Figure 2). Consequently, downstream analyses using these latent variables are also expected to be insensitive to the abundance cutoff used.

### Probabilistic modeling reproduces dataset-specific statistics of microbiomes

To endow generative capacity to our approach, we model the distribution *p*(*z*) of data points over latent variables using Gaussian mixture modeling (**SI Section 3**). Notably, Gaussian mixtures are versatile models to fit multivariate data as they can in principle fit to distributions of arbitrary shapes. From this distribution it is possible to generate realistic *in silico* microbiome compositions by first sampling their latent space embeddings using the learnt distribution and then using the universal species preferences learned during training and obtaining relative species abundances using Eq. 3. Additionally, the Gaussian mixture model also enables to calculate the likelihood of any given latent space coordinate profile relative to the data used to train the model.

Can the generative model sample realistic ecosystem compositions? We tested this using comparison of several lower-order statistics, metrics of distribution density, and embedding accuracy. This is a common approach used in generative models. We trained a generative model using a latent space of dimension *K* = 16 (Figure 2). We tested the accuracy of the generative model by sampling latent variables from an inferred mixture Gaussian *p*(*z*) and then generating *in silico* microbiomes using Eq. 3. For clarity of presentation, we present results for chicken cecum microbiomes. The results for the other datasets are shown in SI Figure 3 and 4. Figure 3A shows that the generative model accurately reproduces OTU mean abundances and standard deviations. Notably, our approach is quite accurate at predicting microbiome composition at all levels of taxonomy (SI Figure 5). Figure 3B (and inset) shows that the model captures species abundance covariation and higher order co-occurrence patterns. Figure 3C shows that the model reproduces the species abundance distribution, including the power law-like scaling between abundances of 10^−3^ and 10^−1^ with a power law exponent of *γ* ∼ − 1.03 (*γ* ∼ − 1.12 for the model) for the inverse cumulative distribution. Figure 3D shows that the model reproduces the distributions of the *α*-diversity metric of Shannon diversity. Figure 3E shows that the model reproduces distribution of the *β*-diversity metric Bray-Curtis dissimilarity between nearest neighbor samples as well as pairs of randomly picked samples.

Are the species preferences towards effective resources *θ*_*ko*_ universal? If so, species preferences inferred from a set of microbiome samples should be able to accurately model the composition of dissimilar microbial communities of the same species. To test this hypothesis, we looked at bacterial microbiomes associated with the cecum and ileum of broiler chickens. The cecum and the ileum are distinct parts of the chicken’s gastrointestinal track. Indeed, the abundances of microorganisms vary substantially between the two organs [42].

We split the cecum microbiome samples randomly into an 80% training and 20% testing split and trained the model only on the training data. As examples of dissimilar ecosystems, we used an additional testing set comprising ileal microbiomes of chickens. As seen in Figure 3F, the species preferences towards effective resources are indeed universal. Unsurprisingly, the learnt species preferences embed testing data from the same organ site (cecum) with similar accuracy as the training data (black versus gray dots, x-axis). Surprisingly, the accuracy of embedding testing data from another organ using the same species preferences was quite similar to testing data from the same organ site (red versus black dots, x-axis). In contrast, the accuracy of embedding the testing data using a scrambled preference matrix was significantly lower (red versus blue dots, x-axis). Similarly, the inferred latent space locations of testing data microbiomes derived from ceca had similar likelihood as the training data. In contrast, the latent space locations of ileal microbiomes had significantly lower likelihood (y-axis). This is consistent with our interpretation of the consumer resource framework. Specifically, while the OTUs have consistent embeddings between the ileum and the cecum, leading to reasonably accurate reconstruction of the ileal microbiomes, the abundances of effective resources and therefore the values of the fitted latent variables *z*s are likely to be markedly different between the two organ sites. Consequently, the latent variables fitted for the ileums have significantly lower likelihood compared to the cecal latent variables. A similar result was observed when comparing embeddings of cattle from the Nordic Red breed using species preferences learnt from Holstein cows (SI Figure 3).

Collectively, these results show that probabilistic modeling of the latent space defined by the C/R model generates microbial compositions that reproduce both universal statistics such as species abundance distributions and dataset specific statistics such as patterns of species covariances and *α*- and *β*-diversity metrics. Moreover, the learnt species-resource preferences are universal; the statistics of probability of observing unseen data and accuracy of embedding unseen data are similar to those of the training data. In other words, the C/R model based generative model can sample compositions of feasible microbiome community structures, an essential feature of accurate generative models.

### Generative modeling identifies host phenotypes that maximally constrain the microbiome

The coarse-grained latent space represents the history of effective resources. Therefore, it can be informative of host phenotypes that affect microbial growth. Indeed, recent work has suggested that many functions of microbial communities are coarse-grainable i.e. explained by a small number of variables [35]. To test this, we investigated the quantitative relationship between phenotypes and associated microbiomes of the rumens of ∼ 800 bovine hosts. Here, ∼ 50 phenotypes were measured, including traits related to rumen chemistry (e.g. volatile fatty acids) and animal physiology (e.g. milk production, feed conversion efficiency) (Supplementary Table 1).

Surprisingly, latent space embeddings inferred only using the microbiome accurately predicted most of the measured phenotypes using linear regression (SI Figure 6). This suggests that the latent variables fitted to the C/R model encode information about the environmental variables experienced by the microbiome. Consequently, these *effective resources* may simultaneously capture microbial community structure and host phenotypes that influence microbial growth. To directly incorporate host phenotypes in our model, we employ a shared latent approach. Specifically, we require the latent variables to simultaneously describe the composition of microbiomes and host phenotypes. To that end, we introduce a new cost function:

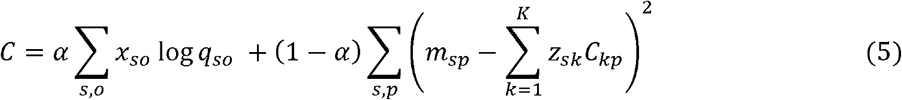

In Eq. 5 the first term captures the goodness of fit to the microbiome composition and the second term is a model to fit the host phenotypes. Unlike the microbiome, the relationship between host phenotypes and latent variables cannot be mechanistically derived. Therefore, we take a data-driven approach. We approximate the host phenotypes using a linear low-rank model (SI Section 1). We note that other functional forms, for example, feed forward neural networks can be employed as well. In Eq. (5), 1 ≥ *α* ≥ 0 decides the relative importance of the two terms in the minimization process. We fit Eq. 7 to the combined microbiome/phenotype data using *K* = 24 latent variables with *α* = 0.975 (chosen to minimize the compromise in accuracy of fitting the microbiome and phenotypes, SI Figure 7). Next, as above, we trained a Gaussian mixture model on the shared latent space (SI Section 3). In this case, sampling latent variables using the learnt distribution *p*(*z*) generated combined microbiome/phenotype data that preserved covariances between the microbiome and the phenotypes (SI Figure 8). Notably, the linear model employed in Eq. 5 may not be sufficient to model host phenotypes from the latent variables. However, at least in this case, nonlinear neural networks did not significantly improve the predictions (SI Figure 6). Therefore, going forward, we employ linear models to connect host phenotypes to latent variables.

A central question in ecology is how strongly environmental parameters or ecosystem functions constrain the community composition. To answer this question in the context of host-associated microbiomes, we used generative modeling. Specifically, we sought to identify which host phenotypes constrain the space of possible microbial community structures? For example, host serum metabolomics may only weakly determine the microbiome while host dietary habits and gut chemistry may have a more constraining impact. Indeed, our model finds that some phenotypes like rumen pH and propionate levels can significantly constrain the space of feasible community structures while others such as organic matter intake do not. Importantly, these dependencies are not always uniform across the range of host phenotypic states. To quantify these relationships, we generated a large number of *in silico* combinations of microbial communities and corresponding host phenotypes. For each phenotype, we evaluated the average Bray-Curtis dissimilarity amongst pairs of communities that corresponded to a narrow range of values of the phenotype. A low dissimilarity amongst communities corresponding to a particular range indicates that specifying the phenotype in that range narrows the space of feasible community compositions. We found that different phenotypes constrained community composition to a different degree. Figures 4ABC show that specifying phenotypes such as levels of rumen short chain fatty acids (propionate and isobutyrate) and rumen pH significantly constrained the microbiome composition in the rumen. In contrast, phenotypes such as organic matter intake and protein intake levels did not constrain the microbiome composition (Supplementary Table 2). Notably, these analyses required *in silico* generation of thousands of feasible microbiome/host phenotype combinations and therefore cannot be performed using the data alone.

**Figure 4.**
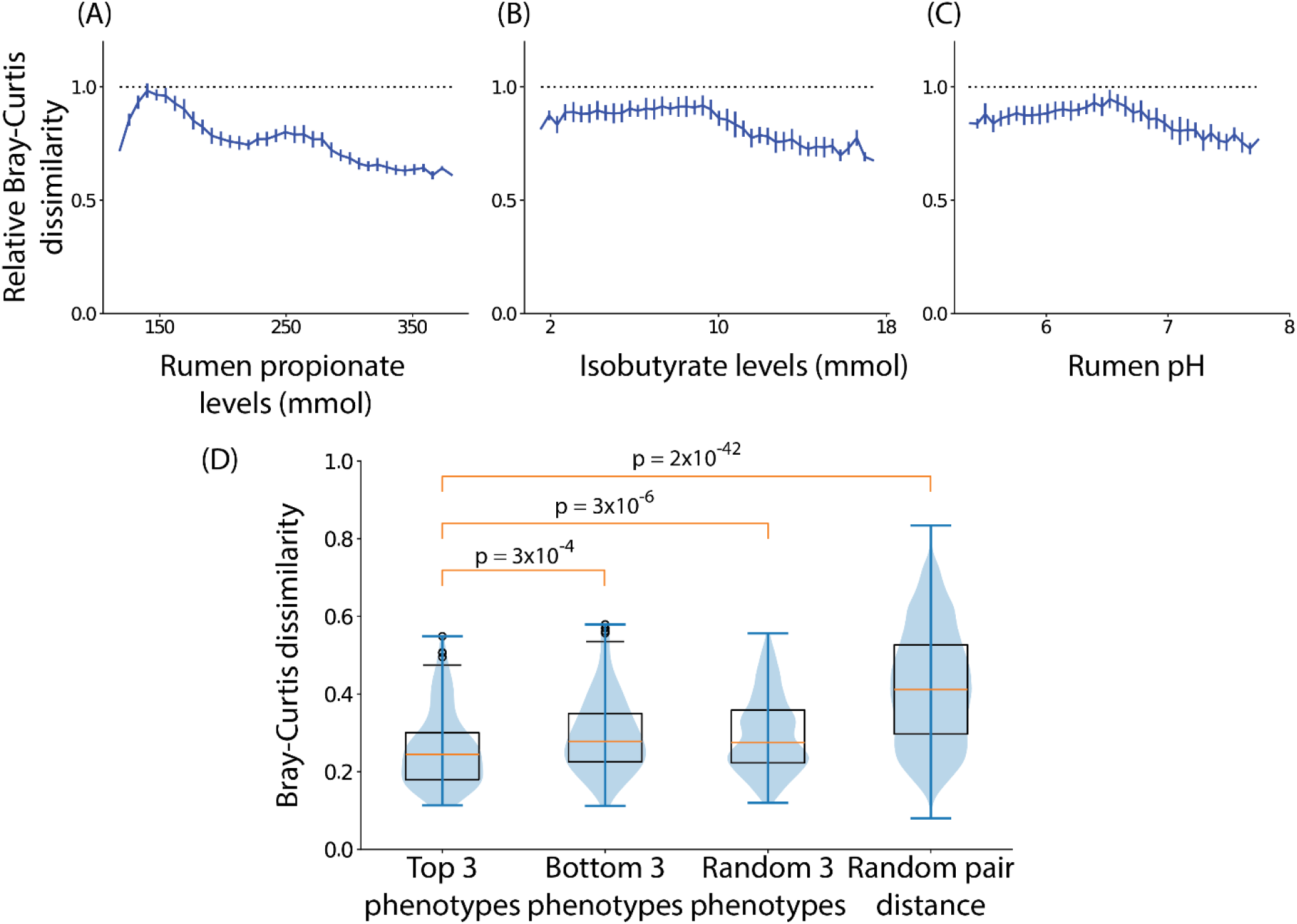
Identifying phenotypes that constrain the microbiome. (A, B, and C) The average Bray-Curtis dissimilarity (y-axis, normalized by the Bray-Curtis dissimilarity between random pairs of communities) amongst *in silico* bovine rumen microbial communities constrained to have a specified single phenotype (x-axis). The top 3 phenotypes that constrained community composition the most are shown. Error bars represent standard errors of the mean. (D) Box plot of Bray-Curtis dissimilarity between predicted microbiome composition and the measured composition using top 3 constraining phenotypes, bottom 3 least constraining phenotypes, 3 randomly chosen phenotypes, and random pairs of communities in the testing samples for bovine rumen communities.

To verify the importance of specific phenotypes in narrowing plausible microbiome compositions, we investigated whether we could reconstruct the microbial composition in the rumens of hosts in the testing set using partial phenotypic information about the host. To do so, we deeply sampled from the mixture gaussian model and found *in silico* samples that had phenotype values in close proximity of the data point of interest. We used the average microbiome composition of these *in silico* microbiomes as our microbiome prediction (SI Section 4). We asked whether partial information about the phenotypes identified in Figure 4ABC could identify the composition of the associated rumen microbiome. Importantly, the predictions made using partial phenotypic information were significantly more accurate compared to the dissimilarity between pairs of randomly picked samples in the testing data.

These results demonstrate that our generative model not only captures complex associations amongst microbial species (Figures 3) but also the associations between the microbiome and corresponding host phenotypes (Figure 4).

### Generative modeling quantifies context-specificity of host-microbiome interactions

Host-microbiome interactions are often context-specific, that is, the association between a taxon and a host phenotype may depend on the composition of the rest of the community. We exploit the generative capacity of our approach to identify subject-specific microbial correlates of methane production (normalized by dry matter intake). Methane production by bovines is a major contributor to greenhouse gasses and microbiome-based therapies to reduce methane production are a promising intervention avenue. To identify host-specific associations, we densely sampled the space of combined microbiome composition and host metadata using our generative model. Next, for each host, we identify *in silico* microbiomes that are in its proximity (evaluated using Bray-Curtis dissimilarity) and evaluate the correlation between microbial OTUs in this subset with methane production. As shown in Figure 5, some OTUs associate with phenotypes in a universal manner, while others associate in a host-specific manner. To illustrate this more clearly, we look at two OTUs classified at the genus level as *Prevotella* and *Moraxellaceae ge* that have similar association strength with methane production when correlated across all samples. In contrast, our model predicts that the *Moraxellaceae ge* OTU correlates with methane across all hosts with similar magnitude while the *Prevotella* OTU has a more heterogeneous association profile.

**Figure 5.**
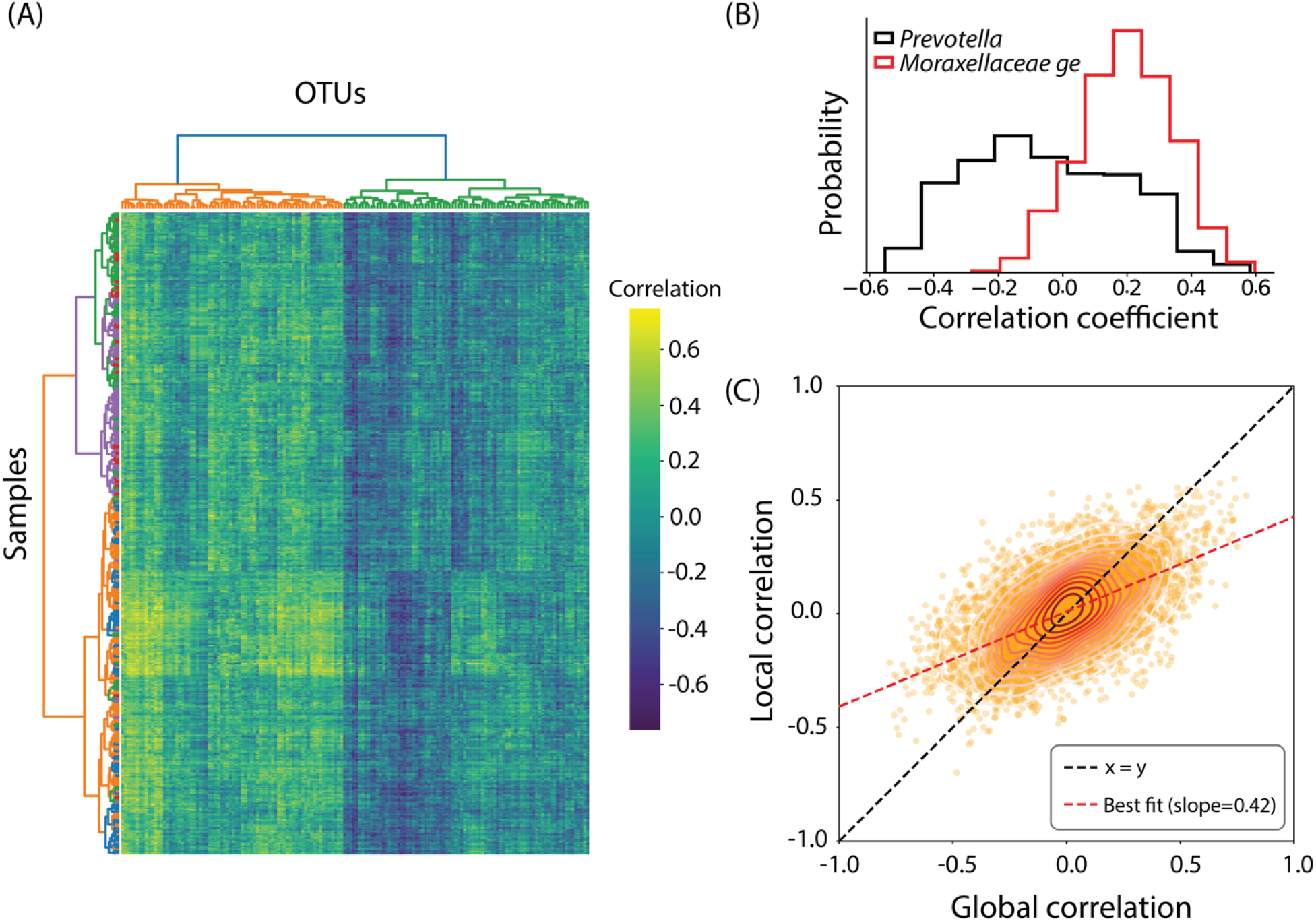
Quantifying context-specificity of host-microbiome associations. (A) Pearson correlation coefficient between methane production (normalized by dry matter intake) and all OTUs evaluated in a sample-specific manner. Different colors in the sample-wise clustering tree represent different farms to which the cows belong. (B) Distribution of sample specific Pearson correlation coefficients for two OTUs (*Prevotella* and *Moraxellaceae ge*). The global (across sample) Pearson correlation for abundance of either OTUs with methane production is ∼ 0.15. (C) The mean host-specific (local) correlation of an OTU with a phenotype is plotted against the global correlation between that OTU and the phenotype, calculated *in silico*. The slope of the best fit line between local and global correlations is 0.42.

The phenotype/OTU associations calculated globally and on a per host basis (averaged over hosts) were correlated with each other. However, the average host-specific correlation was typically half as strong as the global correlation. Notably, some phenotypes were over-represented in host-specific local correlations. For example, of the top 100 host-specific correlated pairs, we found that certain phenotypes occurred more frequently than expected by chance. These include rumen propionate (22 out of 100, hypergeometric test p = 4.08e-16), rumen ammonia (11 out of 100, hypergeometric test p = 1.80e-5), fecal dry matter (11 out of 100, hypergeometric test p = 2.18e-3) and plasma creatinine (6 out of 100, hypergeometric test p = 2.86e-2).

### Designing microbiomes that achieve desired host phenotypes

The chief application of generative models is the design and identification of biological systems that possess user-desired properties, for example, proteins with specific enzymatic activity or structural folds [13]. Here, we illustrate with two examples how our generative modeling can be used to identify host phenotypes and microbial ecosystem compositions that correspond to desired host states.

To validate that our generative model reproduces covariations observed in the data, we investigated the relationship between rumen propionate levels and rumen pH. Propionate production is associated with lower enteric methane emissions in ruminants due to its lower yield of molecular hydrogen [2]. The pH of the rumen is an emergent property arising from the complex acid/base balance of multiple chemicals. Notably, rumen pH negatively correlates with propionate levels (Spearman *r* = − 0.40, *p* = 1.5 × 10^−31^), which was not true of all short chain fatty acids (for example, Spearman *r* = − 0.42, *p* = 30 × 10^−34^ for butyrate and Spearman *r* = 0.10, *p* = 4.5 × 10^−3^ for acetate). Therefore, we sought to characterize host phenotypic states that corresponded with simultaneous high rumen pH and high propionate levels. To achieve this, we performed biased sampling of the latent space using a Markov chain Monte Carlo approach (SI Section 5, red contours in Figure 5A). Notably, these hosts were not observed in the training population (black contours in Figure 5A). Next, we investigated how this shift in propionate and pH changed the levels of other short chain fatty acids. Surprisingly, levels of both acetate and butyrate, which positively correlated with pH, dropped in the biased samples. While not apparent from the data, these results make physiological sense; increasing propionate concentration may be compensated by decrease in acetate and butyrate to maintain mass balance and rumen pH.

Our model can also identify the relationship between host phenotypes and the corresponding composition of the microbiome. Cows fed high amounts of starch often have acidic rumens due to its rapid fermentation compared to fiber rich diets. This may lead to sub-acute ruminal acidosis (SARA), an undesirable condition that leads to lowered economic productivity [46]. Indeed, in the dataset analyzed, starch intake was negatively correlated with pH (Spearman *r* = − 0.53, *p* = 1.65 × 10^−58^). Notably, there were no hosts in the original dataset who could maintain a high pH on a starch-rich diet (black contours in Figure 5C). Given that the pH directly affects microbial growth and can also be controlled by it, we investigated microbial ecosystem compositions that would be present at a high rumen pH in hosts that have a high starch intake using biased sampling of the latent space (**SI Section 5**, red contours in Figure 6C). The rest of the host phenotypes (and the composition of the microbiome) were not specified but automatically sampled according to the biased distribution. The microbiome compositions obtained were significantly different for these desired host phenotypes (Figure 5D). Yet, these *in silico* microbiomes have statistical properties (for example, − *α* and *β* − diversity) that are quite similar to real microbiomes, suggesting that these microbiomes may exist in nature (SI Figure 9). Two OTUs were consistently present in higher abundance in the *in silico* biased samples with high starch intake and a high pH (Supplementary Table 3). These include an OTU from the genus *Ruminobacter* (1 out of 2, hypergeometric test p = 0.025) and an OTU classified as *Succinivibrionaceae_UCG-002*. Both OTUs belonged to the family *Succinivibrionaceae* (2 out of 12, hypergeometric test p = 0.005). Notably, *Ruminobacter* bacteria are known to degrade starch [47]. However, their ability to do so while maintaining a high rumen pH is not fully explored.

**Figure 6.**
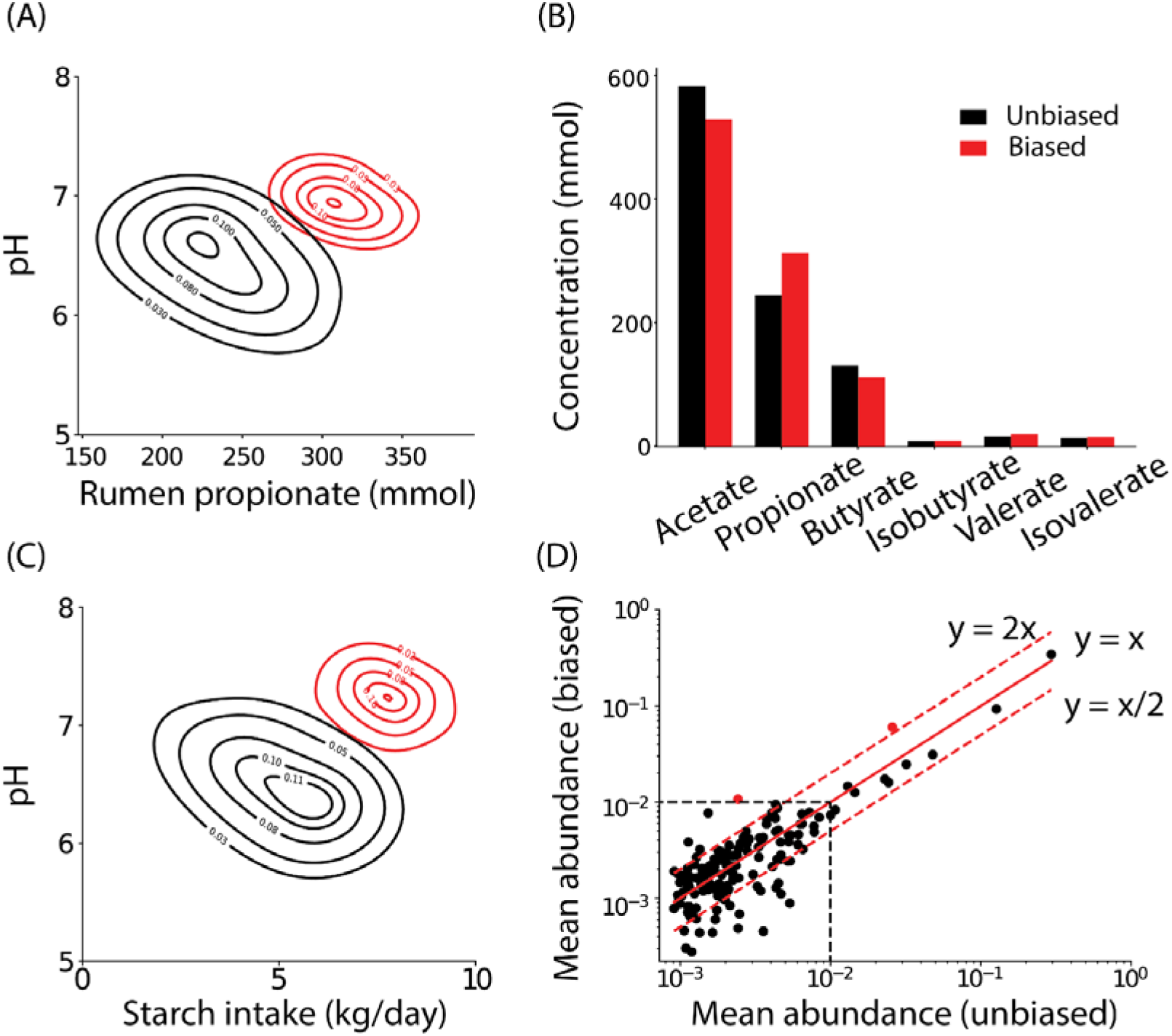
Designing microbiomes for desired phenotypic states. (A) Contour plots of the 2-dimensional histograms of rumen pH and rumen propionate levels. Black contours represent *in silico* communities sampled from the inferred distribution over latent variables. Red contours represent the biased sampling of latent variables which encourages high propionate levels and high rumen pH. (B) Concentrations of volatile fatty acids in unbiased (black) and biased (red) samples. (C) Contour plots of the 2-dimensional histograms of rumen pH and daily starch intake levels. Black contours represent *in silico* communities sampled from the inferred distribution over latent variables. Red contours represent the biased sampling of latent variables which encourages high levels of rumen pH while maintaining a high starch intake. (D) Mean relative abundances of OTUs in the unbiased samples (x-axis) and the biased samples (y-axis). The OTUs with high abundance in the biased samples that also show a significant increase from unbiased samples are colored in red.

These studies show that our model can sample and thereby design host-microbiome metacommunities with user-specified host phenotypes.

## Discussion

Rationalizing the observed variation in complex and high dimensional biological systems using mechanistic modeling is seldom possible. At the same time, we can now collect large amounts of data. This has allowed building data driven generative models that describe possible variations in biological systems. Unfortunately, except for a few examples, studies on host-associated microbiomes usually operate with low sample sizes. Moreover, the context of the microbiome sample (abiotic environment or the phenotypic states of the host organism) is of paramount importance in deciding ecosystem composition. This context dependence likely contributes to a low reproducibility of microbiome studies [47].

In this work, we presented a generative model based on the consumer/resource framework that can model the simultaneous variation in host-associated microbiomes and host’s phenotypes. Notably, our model can assign probabilities to specific communities and phenotypes. This allows us to identify realistic communities that are hypothesized to correspond to desired host states.

Importantly, although our model allows us to sample realistic microbial communities and associated host phenotypes, it does not provide a causal connection between the two. We have shown that certain host traits such as organ pH, nutrient intake, or the concentration of specific metabolites are associated with more specific microbial communities within certain trait value ranges. This does not necessarily mean that those parameters can be used to drive the microbiome towards a particular state, although these are useful hypotheses. Instead, the ability to generate an arbitrary number of realistic *in silico* samples facilitates the study of complex communities by enabling data stratification beyond what is experimentally possible, even to cases that are seldom observed in experimental datasets. As a design tool, the communities obtained by sampling from the model represent target communities for microbiome manipulation strategies including pre and probiotic treatments, antibiotics, microbiome transplants, phage therapies, etc. Knowing what these targets look like should provide clues as to how to achieve them. Notably, however, the model can easily be made causal by incorporating a temporal direction, for example, by training the model on pre- and post-intervention (drugs, probiotics, etc.) microbiomes.

There are several ways to generalize the current framework. *First*, we learnt the latent space embeddings *z*s of individual hosts first and then built a probabilistic model *p*(*z*) that captured the variation in the latent variables. This two-step process can be combined using Expectation-Maximization like algorithms that simultaneously learn embedding for data and model the corresponding mixture Gaussian distribution. *Second*, while the dependence of microbial abundances on latent variables was mechanistically derived, the linear relationship between host phenotypes and latent variables was ad hoc. This linear model can in principle be improved by incorporating more complex relationships between the phenotypes and the latent variables, for example, those modeled using artificial neural networks. *Third*, in the current approach we built a combined latent space model for host phenotypes and the associated microbiomes. Similarly, phenotypic information about microorganisms, for example, consumer/resource preferences inferred using genome-scale metabolic models [48] or species genomic content, can also be incorporated in our framework by requiring the species-specific preferences *θ*_*ko*_ to simultaneously model species abundances in microbiomes and species phenotypes using previously developed latent variable models [17].

While this model was developed for host-associated microbiomes, it can be used to integrate different types of sequencing and phenotypic information not only on animal hosts and corresponding microbial ecosystems but also on cells and their phenotypic properties and potentially in a more general setting when both sequencing and phenotypic information is available for the same entities.

We believe that the framework laid out in the manuscript will be of broad use to microbiome engineers and scientists alike.

## Materials and Methods

All data and code can be found at https://github.com/karthik-yale/host_microbe

### Section 1. Model description and fit to data

#### A. Details of the latent variable model and performance

The consumer/resource (C/R) model [28] has been a popular model in understanding species dynamics in natural and artificial ecosystems. The C/R model imagines that species abundances fluctuate according to their dependence on resources. We have:

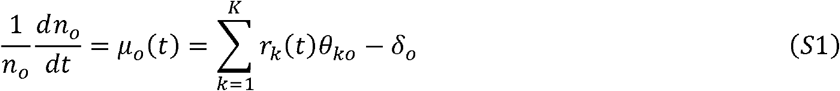

and

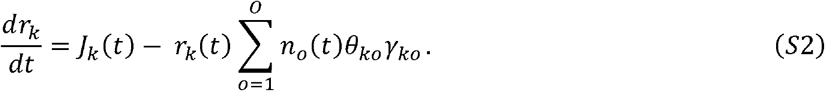

In Eq. S1 and Eq. S2, *n*_*o*_(*t*) ≥ 0 are abundances of consumers (microbial species) “*o*”, *μ*_o_(*t*) is the time-dependent growth rate, *r*_*k*_(*t*) ≥ 0 is the abundance of *resource* “*k*”, *θ*_*ko*_ ≥ 0 are species-preferences for resources, *γ*_*ko*_ ≥ 0 are resource consumption efficiencies, and *J*_*k*_(*t*) are net inflow rates of resources.

With these assumptions, we can formally integrate Eq. 1 to obtain:

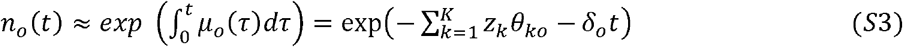

where we recognize

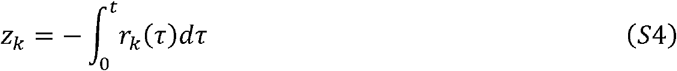

as latent variables that represent the complex temporal dynamics of resources. We note that the death/dilution term can be incorporated into the latent variable representation by identifying *z*_*K*+1_ = *t* (the relevant time scale for the establishment of the current community composition) and *θ*_*K*+1,*o*_ = *δ*_o_. Shifting the indices from *K* + 1 to *K* for simplicity, the relative abundances of organisms *q*_*o*_ are simple functions of the latent variables:

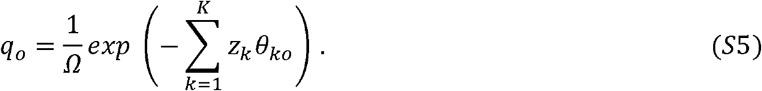

Eq. S5 can be used either in a longitudinal context where latent variables *z*_*k*_ evolve over time inside a single host or in a cross-sectional context where we imagine that microbiomes associated with similar hosts (for example, same host organism) couple to the latent variables in the same manner and that latent variables represent snapshots of individual hosts.

Here, we describe in detail the most general formulation of our approach; to model a collection of host-associated ecosystems along with the phenotypes of the host. We start with the measurement of relative abundances of species (or operational taxonomic units, OTUs) in microbial ecosystems associated with hosts. We denote these abundances as *x*_*so*_ where *S* ∈ [1, *S*] is the index of the subject, *o*” is an organism. Additionally, we assume that measurements of *P* phenotypic metadata *m*_*sp*_(*p* ∈ [1, *P*]) that can potentially affect the microbiome are also measured.

#### B. Fitting the model to microbiome data

We write the model-predicted composition of the ecosystems as

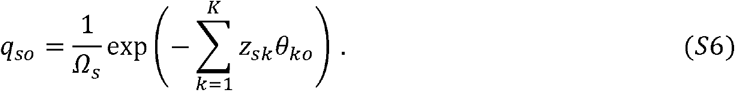

Since the latent variables are related to the properties of the ecosystem and the host, we further hypothesize that the host’s phenotypic metadata explain the latent variables. Unfortunately, however, the relationship between latent variables and host phenotypes cannot be derived using mechanistic models. To that end, we use data-driven methods to capture this dependence. Specifically, we use a generalized low rank linear model:

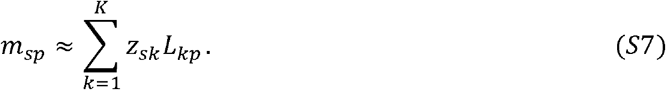

where *g*(·) is a user-specified function. We use *g*(·) = *I*(·) (identity function). Importantly, we require that the same host-specific latent variables explain the composition of all ecosystems as well as the host’s phenotypic information. To that end, we write a combined cost function:

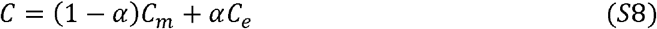

where for the bovine rumen data, *C*_*m*_ is the squared *L*_2_ loss between the host metadata and the corresponding model prediction. *C*_*e*_ is the Kullback-Leibler divergence between the measured microbial composition in the ecosystem *and* the corresponding model prediction. 0 ≤ *α* ≤ 1 is a scaling factor that determines the relative importance of the two terms. When fitting only the microbiome data, we set *α* =0. We have the derivatives:

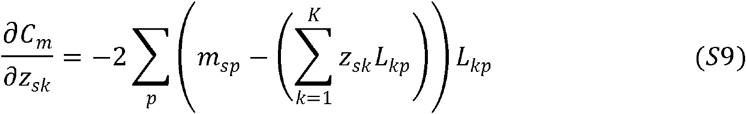

and

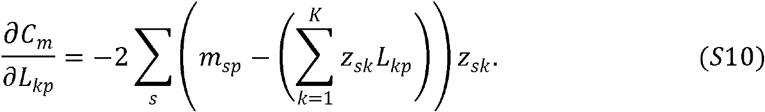

The derivatives for the cost *C*_*e*_ are

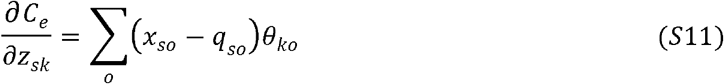

and

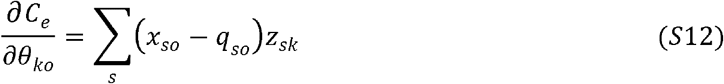

We infer the model parameters using gradients in Eq. S9-S12 and constant step-size gradient descent with learning rate *η* ranging between *η* =10^−3^ −10^−5^ depending on the dataset. The learning rate was set to ensure that the loss decreased smoothly with the number of iterations. We stopped the learning process when the sum of *L*_2_ norms of the gradients divided by *L*_2_ norms of the corresponding matrices was less than 10^−2^. Notably, the inference problem is convex in ***Z***s, ***θ***s, and ***L*** when the other parameters are kept constant [49]. Therefore, the inferred matrices were robust (up to a rotation) to hyperparameters related to learning.

### Section 2. Details of the datasets

Three datasets were used in our study that spanned multiple animal hosts. We used data collected on (1) bacterial microbiome compositions in the bovine rumen as well as physiological information about the bovine host, (2) bacterial ecosystems in chicken ceca, (3) fecal bacterial microbiomes in humans. Below, we provide details of individual data sets.

#### Bovine rumen microbiome and host physiology

The data on rumen microbiomes and host physiology of Holstein cows was downloaded from the paper by Wallace et al. [37]. For microbiome profiling, raw sequencing reads targeting the bacterial 16S rRNA gene were downloaded from SRA, and processed with DADA2 (v.1.12.1) [50] to generate a matrix of read counts per sample at the level of amplicon sequence variants (ASVs). ASVs were clustered into 97% identity OTUs using CD-HIT [51] and OTU-level relative abundances were calculated from the ASV counts using the corresponding clusters. The *assignTaxonomy* method of DADA2 was used to assign genus-level taxonomic labels to the OTUs based on the Silva v. 138 database [52]. We only included bacterial OTUs whose average relative abundance was larger than 0.1%. This cutoff was based on our previous work where we had shown that OTU abundance variation below 0.1% is likely to be dominated by technical noise. In the end, we had data on relative abundances of 156 bacterial OTUs in 790 cows. An additional OTU represented the combined abundance of all OTUs whose average abundance was below 0.1%. For training/testing purposes, the data was randomly split into a 80% training set and a 20% testing set. The OTUs identified in Holstein cows were also used to define the microbiomes of Nordic red cows. There were 156 bovine hosts belonging to the Nordic red species.

#### Chicken microbiomes

The data on chicken cecal microbiomes was downloaded from the paper by Johnson et al. [38]. Raw 16S rRNA amplicon sequences were processed as described above. As above, we only included OTUs whose average relative abundance was larger than 0.1%. In the end, we had a total of 425 host samples and 147 OTUs. As above, one OTU represented the combined abundance of all OTUs whose average abundance was below 0.1%.

#### Human fecal microbiome

Metagenomics-derived genus-level microbial abundances were obtained from the Inflammatory Bowel Disease database [39]. We considered 127 samples from the healthy controls. As above, genera with average relative abundances across samples lower than 0.1% were aggregated into a single taxon, yielding a total of 95 bacterial genera analyzed.

### Section 3. Mixture Gaussian models

In our model, the compositions of the microbiomes and host metadata are direct functions of the latent space variables. We make our model generative by approximating a distribution over the latent space. To that end, we fit a mixture multivariate Gaussian model using Python’s sklearn.mixture.GaussianMixture to the inferred latent space descriptors. The number of Gaussians in the mixture model is varied between 1 and 5 and the mixture with the lowest Bayesian information criterion (BIC). Given that fitting a mixture Gaussian model typically leads to a locally optimum solution, the number of Gaussians in the mixture and the mean values and covariances matrices of the mixtures may change from fit to fit. Therefore, we ran the procedure 100 times and chose the Gaussian mixture model with the lowest BIC.

### Section 4. Predicting microbiome composition from partial host phenotypic data

To identify microbiome compositions corresponding to specified partial list of host phenotypes, we deeply sampled the inferred mixture Gaussian distribution and retained the samples with phenotypes of interest with values in proximity to the host values (within two tenths of the standard deviations of the measured phenotypes). We use the mean composition of the communities identified to have the required phenotype values as our predicted microbiome composition. The Bray-Curtis dissimilarity between this predicted microbiome composition and the host microbiome composition was used as the error/distance measure.

### Section 5. Biased sampling of the latent space to identify microbiome compositions corresponding to desired host states

To identify microbiome compositions corresponding to desired host phenotypes, we sampled latent variables using the inferred mixture Gaussian distribution amended with a biasing term. Specifically, we modified the log probability of latent variables as

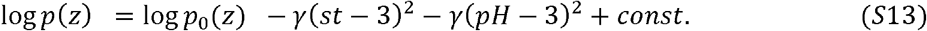

In Eq. S13, *p*_0_(*z*) is the inferred mixture Gaussian model, *st* denotes z-scored starch intake and *pH* denotes z-scored rumen pH. The starch intake and the pH were predicted from the latent variables using the inferred linear model (Eq. S7). *γ* = 0 corresponded to the unbiased simulation and biasing was achieved by setting *γ* = 2. To sample the latent space, we randomly chose 100 starting points from *p*_0_(*z*) and performed a Markov chain Monte Carlo simulation with the modified probability. Once the latent variables are sampled, microbiome compositions are predicted using Eq. S6.

## Acknowledgments

PD is supported by NIGMS grant R35GM142547. PD and KS are supported by the Bill and Melinda Gates Foundation grant INV-062437 to BiomEdit.

## Notes

### Competing Interest Statement

GP is an employee and owns profit interest in BiomEdit LLC.

### Summary of Updates

Significantly extended the analysis with a couple of new figures on context-specific host-microbiome associations.

## References

1. Gilbert, J.A., et al., Current understanding of the human microbiome. Nat Med, 2018. 24(4): p. 392–400.

2. Tapio, I., et al., The ruminal microbiome associated with methane emissions from ruminant livestock. J Anim Sci Biotechnol, 2017. 8: p. 7.

3. Grilli, J., Macroecological laws describe variation and diversity in microbial communities. Nat Commun, 2020. 11(1): p. 4743.

4. Ji, B.W., et al., Macroecological dynamics of gut microbiota. Nat Microbiol, 2020. 5(5): p. 768–775.

5. Ji, B.W., et al., Quantifying spatiotemporal variability and noise in absolute microbiota abundances using replicate sampling. Nat Methods, 2019. 16(8): p. 731–736.

6. Turnbaugh, P.J., et al., The human microbiome project. Nature, 2007. 449(7164): p. 804–10.

7. Marti, J.M., et al., Health and Disease Imprinted in the Time Variability of the Human Microbiome. mSystems, 2017. 2(2).

8. Obeng, N., et al., Evolution of Microbiota-Host Associations: The Microbe’s Perspective. Trends Microbiol, 2021. 29(9): p. 779–787.

9. Bansept, F., et al., Modeling host-associating microbes under selection. ISME J, 2021. 15(12): p. 3648–3656.

10. Levy, R. and E. Borenstein, Metabolic modeling of species interaction in the human microbiome elucidates community-level assembly rules. Proc Natl Acad Sci U S A, 2013. 110(31): p. 12804–9.

11. Kinross, J.M., A.W. Darzi, and J.K. Nicholson, Gut microbiome-host interactions in health and disease. Genome Med, 2011. 3(3): p. 14.

12. Lopez, R., A. Gayoso, and N. Yosef, Enhancing scientific discoveries in molecular biology with deep generative models. Mol Syst Biol, 2020. 16(9): p. e9198.

13. Wu, Z., et al., Protein sequence design with deep generative models. Curr Opin Chem Biol, 2021. 65: p. 18–27.

14. Sgarbossa, D., U. Lupo, and A.F. Bitbol, Generative power of a protein language model trained on multiple sequence alignments. Elife, 2023. 12.

15. Akl, H., et al., GENERALIST: A latent space based generative model for protein sequence families. PLoS Comput Biol, 2023. 19(11): p. e1011655.

16. Yelmen, B., et al., Creating artificial human genomes using generative neural networks. PLoS Genet, 2021. 17(2): p. e1009303.

17. Zhao, X., G. Plata, and P.D. Dixit, SiGMoiD: A super-statistical generative model for binary data. PLoS Comput Biol, 2021. 17(8): p. e1009275.

18. Xu, C., et al., Probabilistic harmonization and annotation of single-cell transcriptomics data with deep generative models. Mol Syst Biol, 2021. 17(1): p. e9620.

19. Ekeberg, M., et al., Improved contact prediction in proteins: using pseudolikelihoods to infer Potts models. Phys Rev E Stat Nonlin Soft Matter Phys, 2013. 87(1): p. 012707.

20. Frazer, J., et al., Disease variant prediction with deep generative models of evolutionary data. Nature, 2021. 599(7883): p. 91–95.

21. Rong, R., et al., MB-GAN: Microbiome Simulation via Generative Adversarial Network. Gigascience, 2021. 10(2).

22. Reiman D, D.Y., Using Conditional Generative Adversarial Networks to Boost the Performance of Machine Learning in Microbiome Datasets. biorXiv, 2020.

23. Choi, J.M., et al., DeepMicroGen: a generative adversarial network-based method for longitudinal microbiome data imputation. Bioinformatics, 2023. 39(5).

24. Oh, M. and L. Zhang, Generalizing predictions to unseen sequencing profiles via deep generative models. Sci Rep, 2022. 12(1): p. 7151.

25. Menon, R., V. Ramanan, and K.S. Korolev, Interactions between species introduce spurious associations in microbiome studies. PLoS Comput Biol, 2018. 14(1): p. e1005939.

26. Holmes, I., K. Harris, and C. Quince, Dirichlet multinomial mixtures: generative models for microbial metagenomics. PLoS One, 2012. 7(2): p. e30126.

27. Ho, P.Y., B.H. Good, and K.C. Huang, Competition for fluctuating resources reproduces statistics of species abundance over time across wide-ranging microbiotas. Elife, 2022. 11.

28. Marsland, R., 3rd, W. Cui, and P. Mehta, A minimal model for microbial biodiversity can reproduce experimentally observed ecological patterns. Sci Rep, 2020. 10(1): p. 3308.

29. Guthrie, L., et al., Impact of a 7-day homogeneous diet on interpersonal variation in human gut microbiomes and metabolomes. Cell Host Microbe, 2022. 30(6): p. 863–874 e4.

30. Chesson, P., MacArthur’s consumer-resource model. J Th Biol, 1990. 37(1): p. 26–38.

31. Goyal, A., et al., Ecology-guided prediction of cross-feeding interactions in the human gut microbiome. Nat Commun, 2021. 12(1): p. 1335.

32. Dubinkina, V., et al., Multistability and regime shifts in microbial communities explained by competition for essential nutrients. Elife, 2019. 8.

33. Goldford, J.E., et al., Emergent simplicity in microbial community assembly. Science, 2018. 361(6401): p. 469–474.

34. M, M.J.a.T., Emergent predictability in microbial ecosystems. biorXiv, 2024.

35. M, M.J.a.T., Defining Coarse-Grainability in a Model of Structured Microbial Ecosystems. Physical Review X, 2022. 12: p. 021038.

36. Ho PY, N.T., Sanchez JM, DeFelice BC, Huang KC, Resource competition predicts assembly of in vitro gut bacterial communities. biorXiv, 2022.

37. Dal Bello, M., et al., Resource-diversity relationships in bacterial communities reflect the network structure of microbial metabolism. Nat Ecol Evol, 2021. 5(10): p. 1424–1434.

38. Thompson, L.R., et al., A communal catalogue reveals Earth’s multiscale microbial diversity. Nature, 2017. 551(7681): p. 457–463.

39. Hart, S.F.M., et al., Uncovering and resolving challenges of quantitative modeling in a simplified community of interacting cells. PLoS Biol, 2019. 17(2): p. e3000135.

40. Shahin, M., B. Ji, and P.D. Dixit, EMBED: Essential MicroBiomE Dynamics, a dimensionality reduction approach for longitudinal microbiome studies. NPJ Syst Biol Appl, 2023. 9(1): p. 26.

41. Wallace, R.J., et al., A heritable subset of the core rumen microbiome dictates dairy cow productivity and emissions. Sci Adv, 2019. 5(7): p. eaav8391.

42. Johnson, T.J., et al., A Consistent and Predictable Commercial Broiler Chicken Bacterial Microbiota in Antibiotic-Free Production Displays Strong Correlations with Performance. Appl Environ Microbiol, 2018. 84(12).

43. Lloyd-Price, J., et al., Multi-omics of the gut microbial ecosystem in inflammatory bowel diseases. Nature, 2019. 569(7758): p. 655–662.

44. Fisher, C.K., T. Mora, and A.M. Walczak, Variable habitat conditions drive species covariation in the human microbiota. PLoS Comput Biol, 2017. 13(4): p. e1005435.

45. Beasley, D.E., et al., The Evolution of Stomach Acidity and Its Relevance to the Human Microbiome. PLoS One, 2015. 10(7): p. e0134116.

46. Kleen, J.L., et al., Subacute ruminal acidosis (SARA): a review. J Vet Med A Physiol Pathol Clin Med, 2003. 50(8): p. 406–14.

47. Schloss, P.D., Identifying and Overcoming Threats to Reproducibility, Replicability, Robustness, and Generalizability in Microbiome Research. mBio, 2018. 9(3).

48. Plata, G., C.S. Henry, and D. Vitkup, Long-term phenotypic evolution of bacteria. Nature, 2015. 517(7534): p. 369–72.

49. Dixit, P.D., Thermodynamic inference of data manifolds. Physical Review Research, 2020. 2: p. 023201.

50. Callahan, B.J., et al., DADA2: High-resolution sample inference from Illumina amplicon data. Nat Methods, 2016. 13(7): p. 581–3.

51. Li, W. and A. Godzik, Cd-hit: a fast program for clustering and comparing large sets of protein or nucleotide sequences. Bioinformatics, 2006. 22(13): p. 1658–9.

52. Yilmaz, P., et al., The SILVA and “All-species Living Tree Project (LTP)” taxonomic frameworks. Nucleic Acids Res, 2014. 42(Database issue): p. D643–8.

