## Supplementary Figures for "Designing host-associated microbiomes using the consumer/resource model"

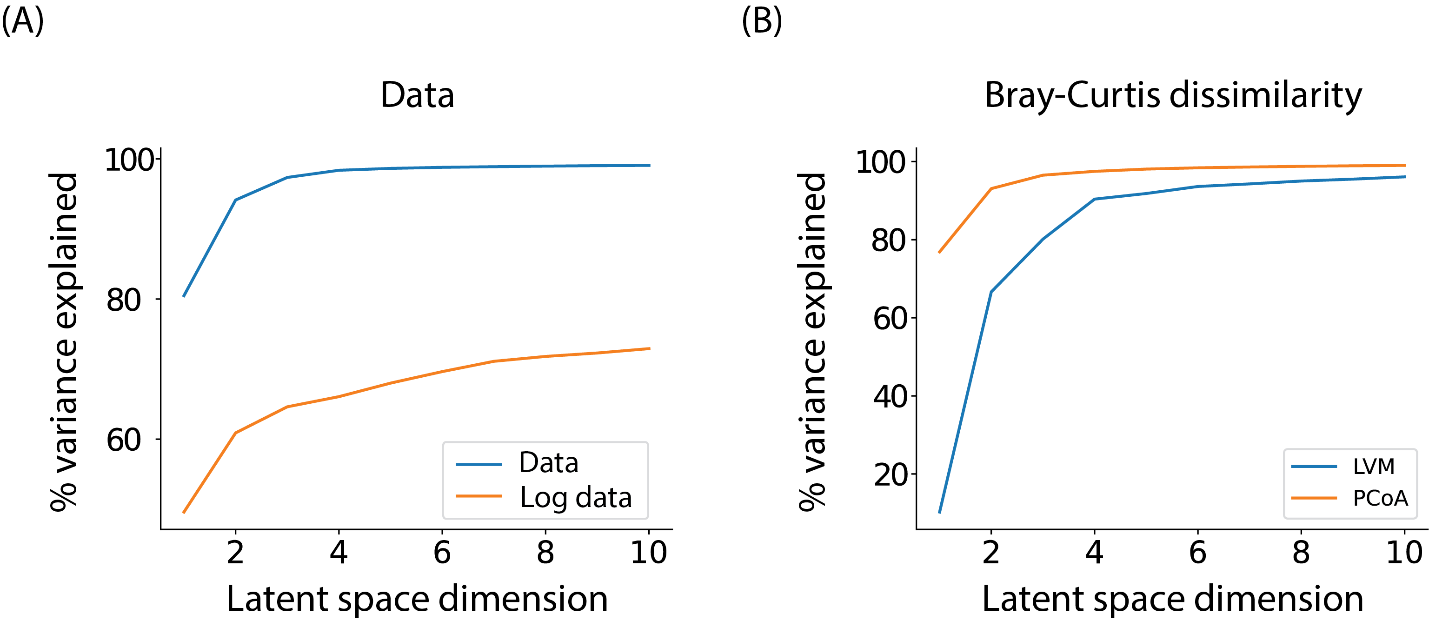

**SI figure 1. Latent variable model fits accurately to data.** (A) The squared Pearson correlation coefficient between the abundance data matrix and log of the abundance data matrix and their corresponding model reconstruction as a function of $K,$ the number of latent variables. (B) The squared Pearson correlation coefficient between the Bray-Curtis dissimilarity (BCD) matrix and the BCD matrix reconstructed using the latent variable model as well as PCoA.

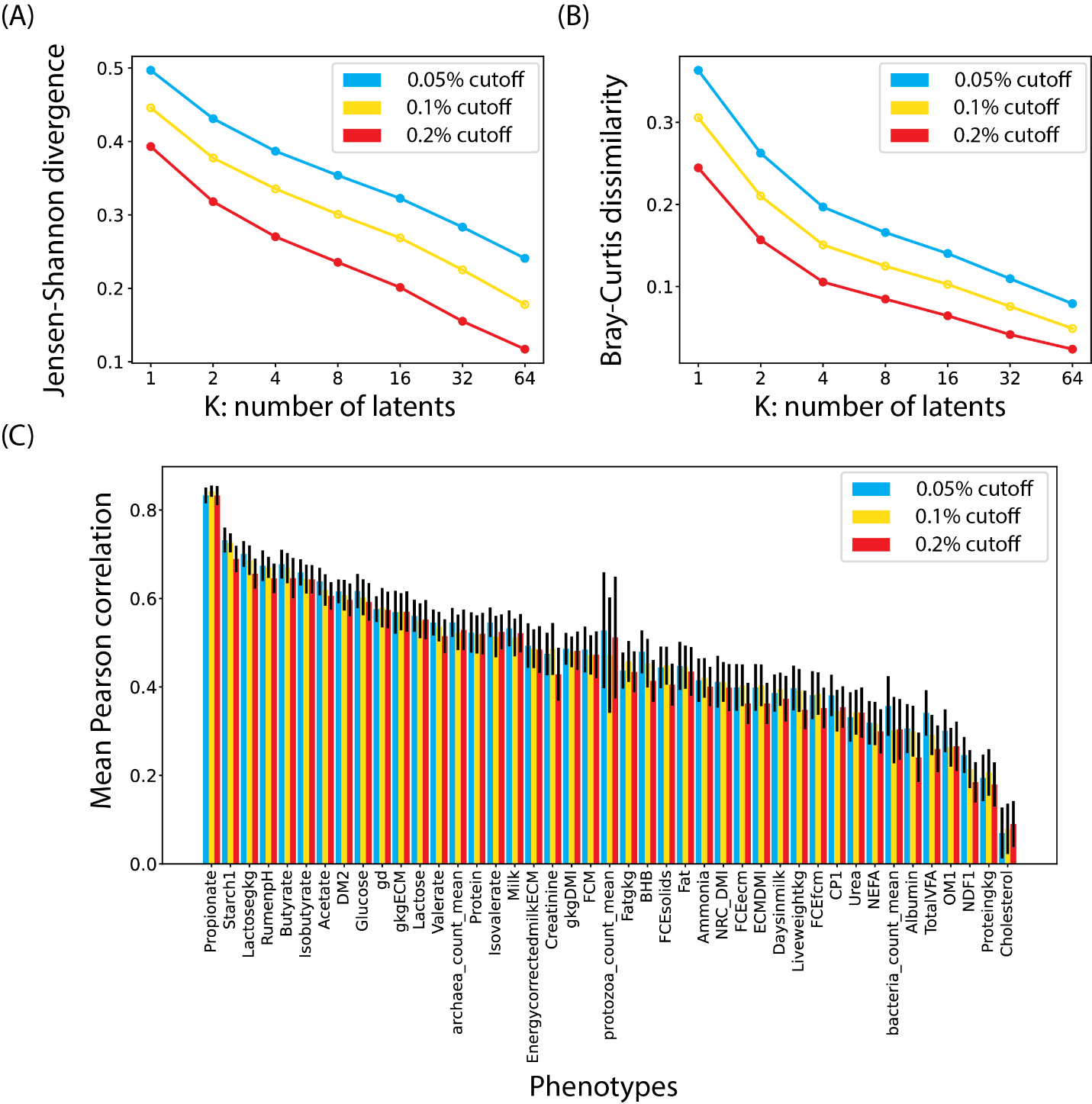

**SI figure 2.** Panels (A) and (B) show the average Jensen-Shannon distance and the average Bray-Curtis dissimilarity between observed community composition and the corresponding model fit as a function of the dimension of the latent space respectively for datasets processed with different cutoffs on mean abundances. The exponential decrease in the error metrics illustrates that the low dimensionality of microbiomes is not dependent on the cutoffs. (C) A bar graph of Pearson correlation coefficients for host metadata predicted from latent variables learnt solely from microbiomes. Pearson correlation coefficients were computed using a simple linear model. The test-train split of test size 25%, was repeated 30 times and the mean values of the Pearson correlation coefficients are plotted with the error bars signifying the standard deviation in the Pearson correlation coefficients. The mean Pearson correlation does not vary significantly across different cutoffs on the mean abundance.

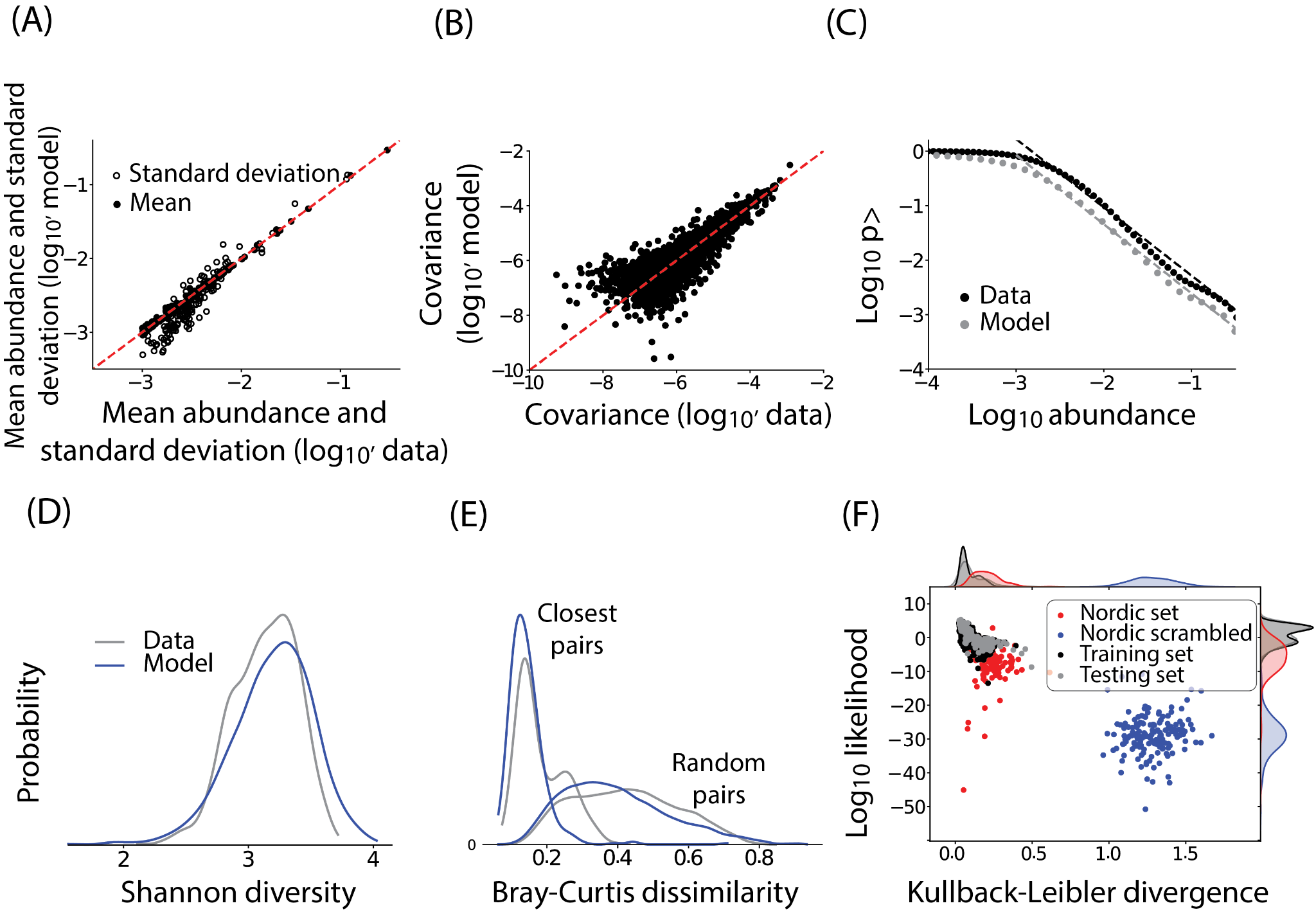

**SI figure 3. Latent space-based model generate accurate *in silico* rumen microbiome communities.** (A) A comparison between mean OTU abundances (filled circles) and standard deviations (open circles) and the corresponding model predictions (y-axis). The dashed red line represents x=y. (B) A comparison between OTU-OTU covariance as computed in the data and as predicted by the model. Absolute values are shown. (C) The inverse cumulative distribution representing the probability of observing an OTU abundance greater than a given abundance (x-axis) as computed from the data (black) and the corresponding model prediction (grey). The dashed lines show the best fit power law between relative abundances of ${10}^{-3}$ and ${10}^{-1}$. (D) The distribution of Shannon diversity of community compositions as observed in the data (gray) and computed from the model generated communities (blue). (E) The distribution of Bray-Curtis dissimilarity between random pairs of community as well as closest pairs of communities in the data (gray) and the model (blue). Distributions are smoothed using Gaussian kernel density estimation. (F) a scatter plot of Kullback-Leibler divergences between community composition in the data and the corresponding model fit (x-axis) and the log-likelihood of the embedded latent variables (y-axis). Data are shown for the training data, testing data, validation data and validation data (Nordic cows) learned with a scrambled preference matrix.

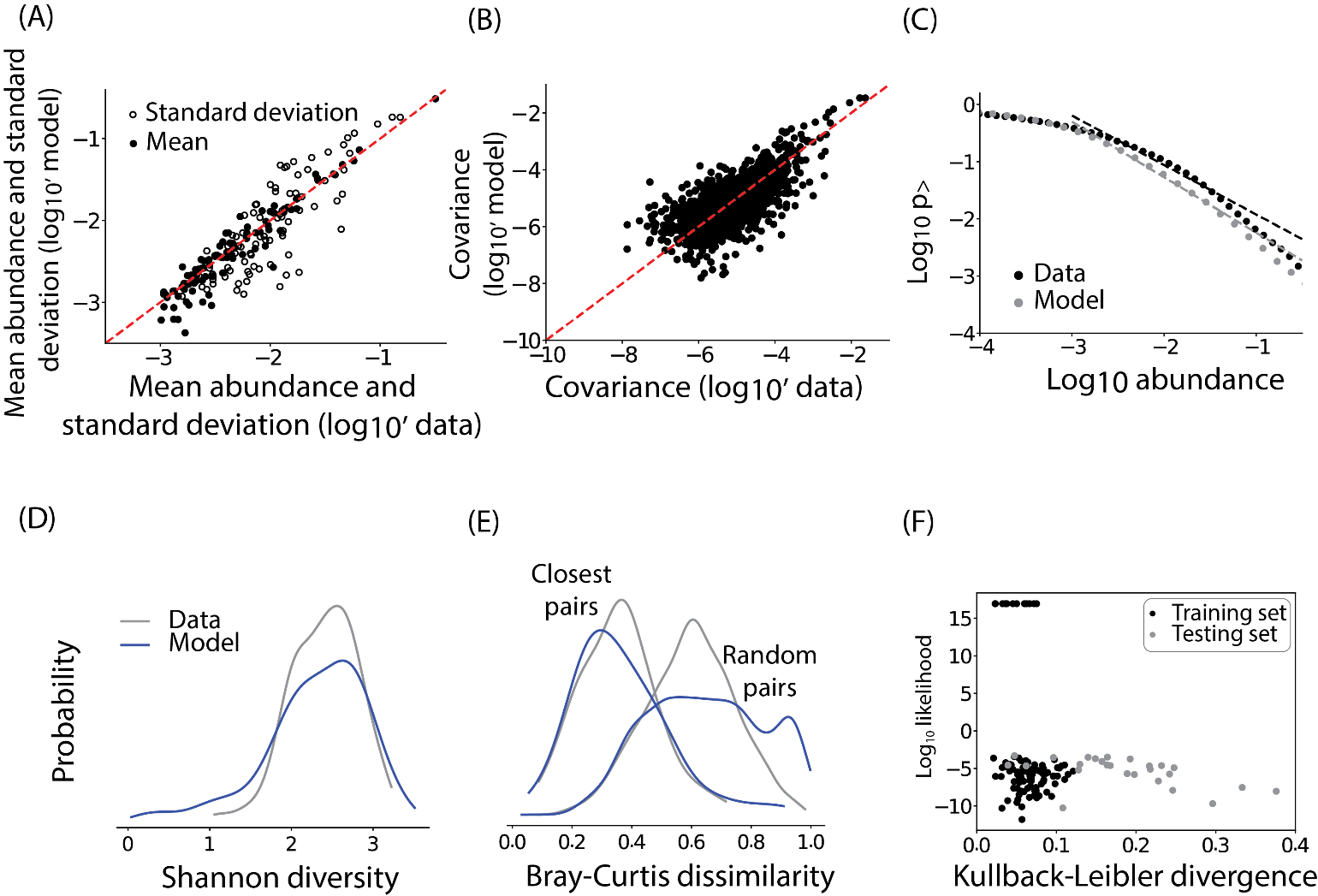

**SI figure 4. Latent space-based model generate accurate *in silico* communities (Human)** (A) A comparison between mean OTU abundances (filled circles) and standard deviations (open circles) and the corresponding model predictions (y-axis). The dashed red line represents x=y. (B) A comparison between OTU-OTU covariance as computed in the data and as predicted by the model. Absolute values are shown. (C) The inverse cumulative distribution representing the probability of observing an OTU abundance greater than a given abundance (x-axis) as computed from the data (black) and the corresponding model prediction (grey). The dashed lines show the best fit power law between relative abundances of ${10}^{-3}$ and ${10}^{-1}$. (D) The distribution of Shannon diversity of community compositions as observed in the data (gray) and computed from the model generated communities (blue). (E) The distribution of Bray-Curtis dissimilarity between random pairs of community as well as closest pairs of communities in the data (gray) and the model (blue). Distributions are smoothed using Gaussian kernel density estimation. (F) a scatter plot of Kullback-Leibler divergences between community composition in the data and the corresponding model fit (x-axis) and the log-likelihood of the embedded latent variables (y-axis). Data are shown for the training (black) and testing data (grey).

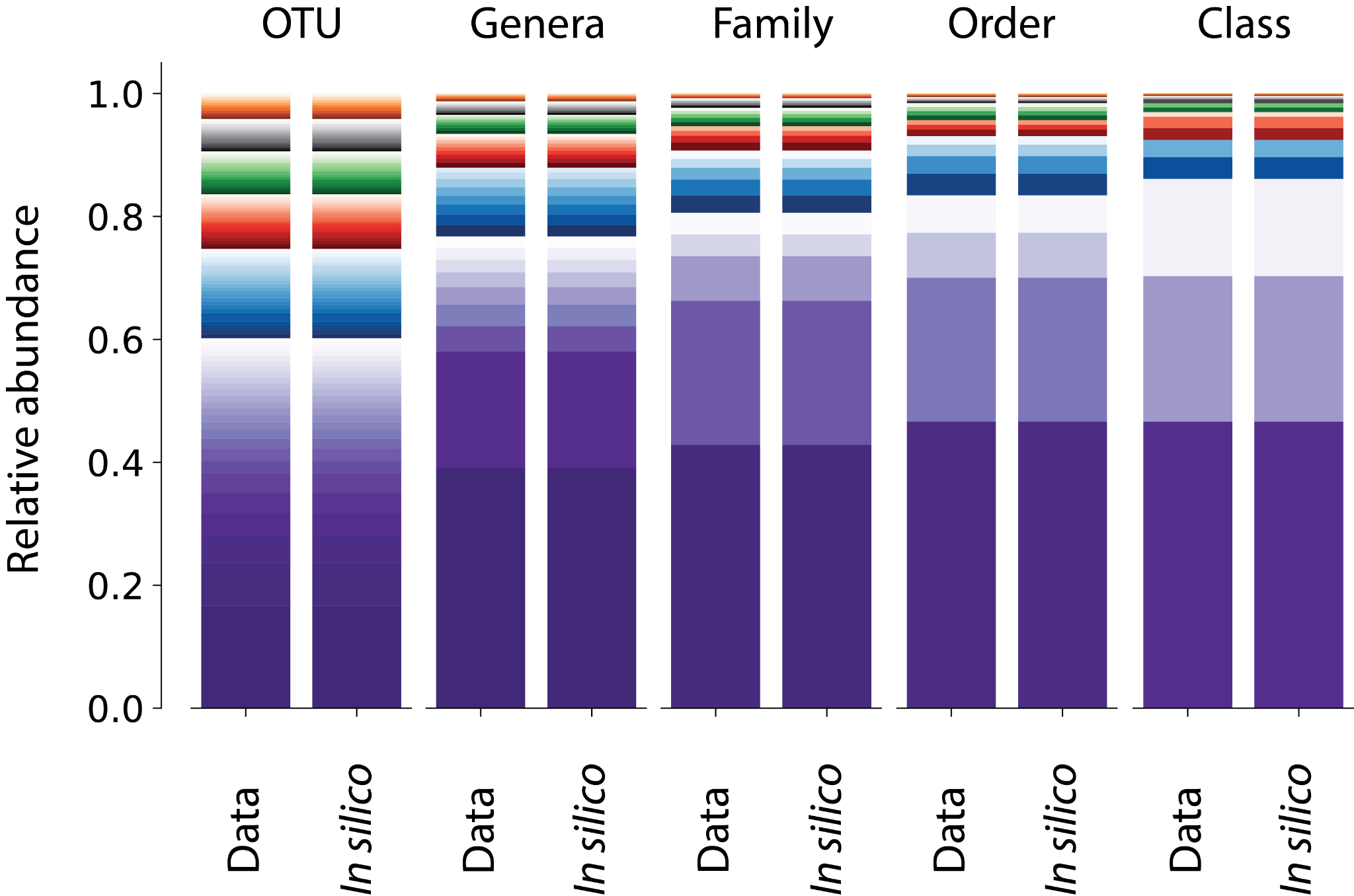

**SI figure 5.** Generative model is accurate at predicting microbiome composition at all levels of taxonomy. The mean abundance of OTUs at different levels of taxonomical classification are plotted for data and *in silico* communities.

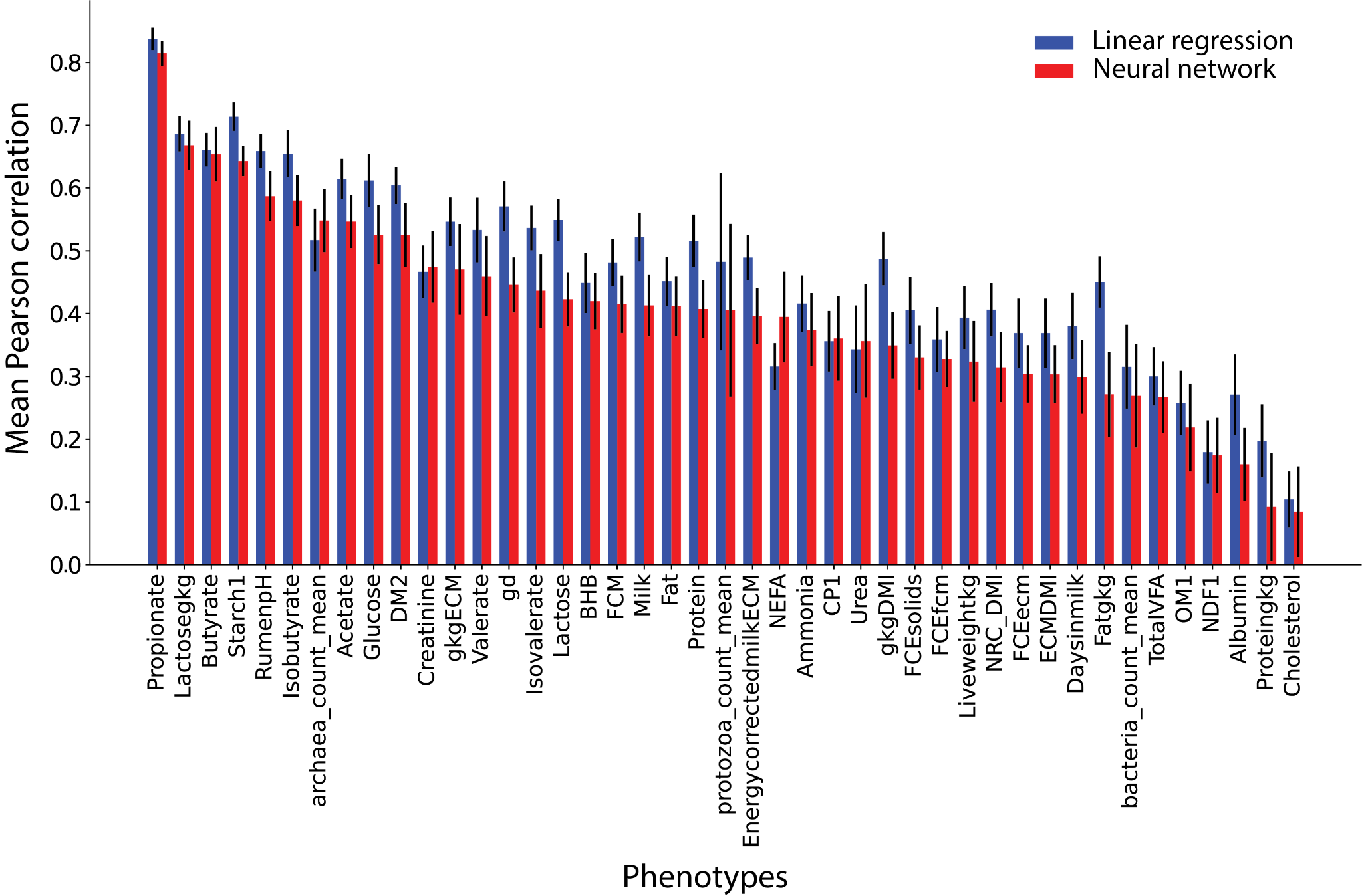

**SI figure 6.** A bar graph of Pearson correlation coefficients for out of sample host metadata predicted from latent variables learnt solely from microbiomes. Pearson correlation coefficients were computed using a simple linear model (blue) (as is done in the main text) as well as a single layer neural network with a sigmoid activation function (red). The test-train split of test size 25%, was repeated 30 times and the mean values of the Pearson correlation coefficients are plotted with the error bars signifying the standard deviation in the Pearson correlation coefficients.

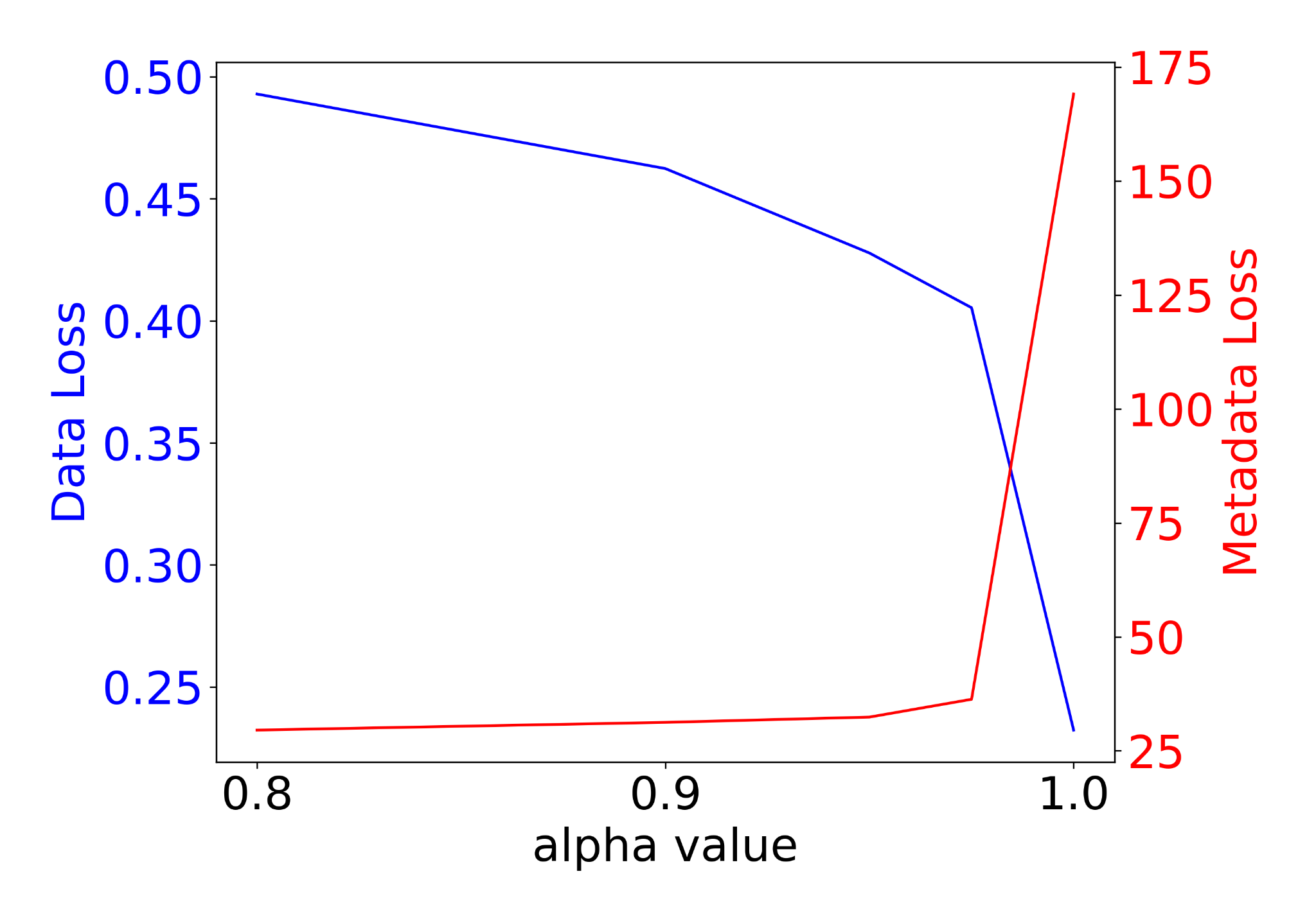

**SI figure 7.** Alpha value was chosen to minimize both phenotype and community fit error.

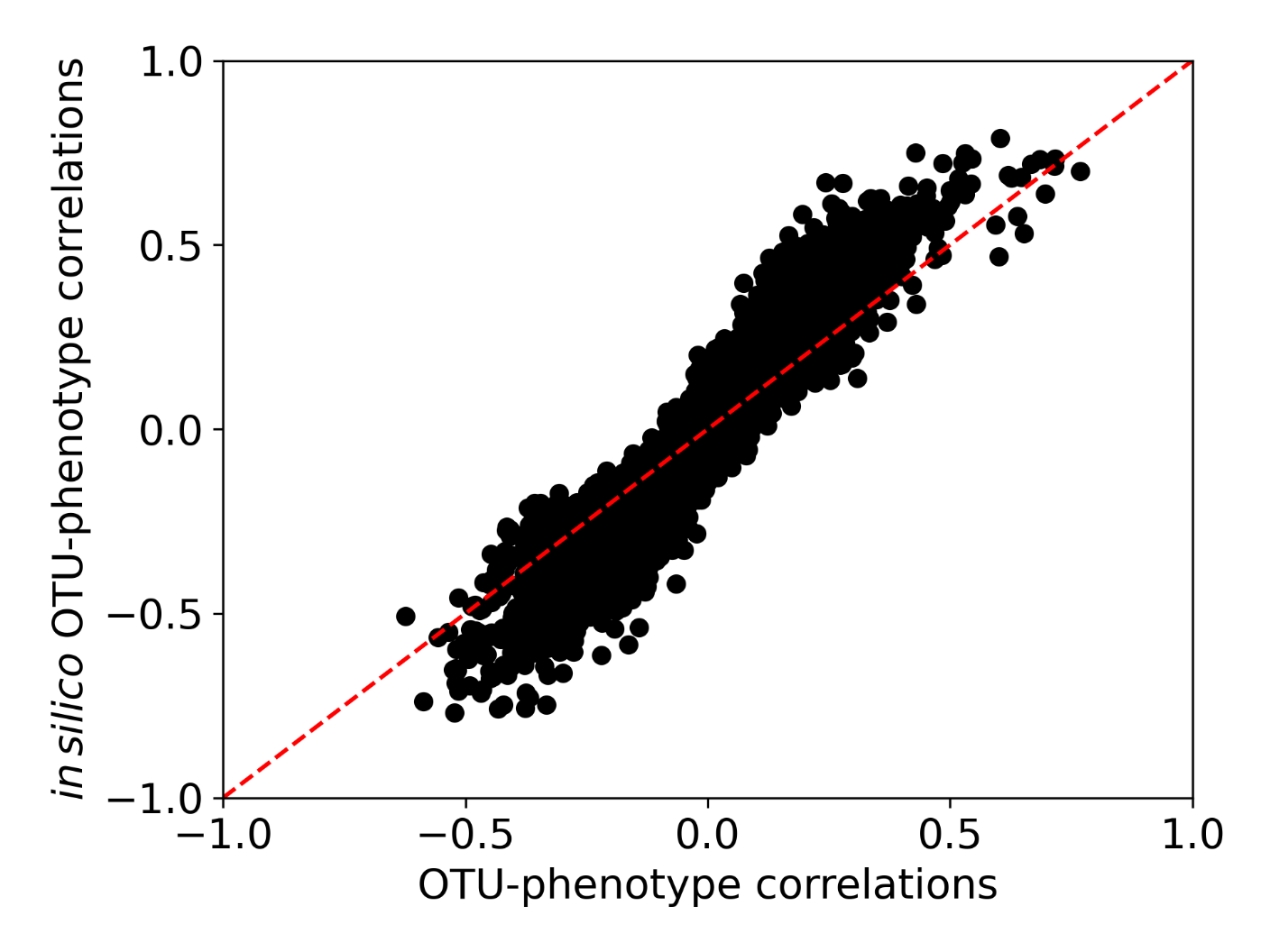

**SI figure 8.** Generative model on simultaneous latent space representation of both communities and their metadata captures OTU-phenotype correlations.

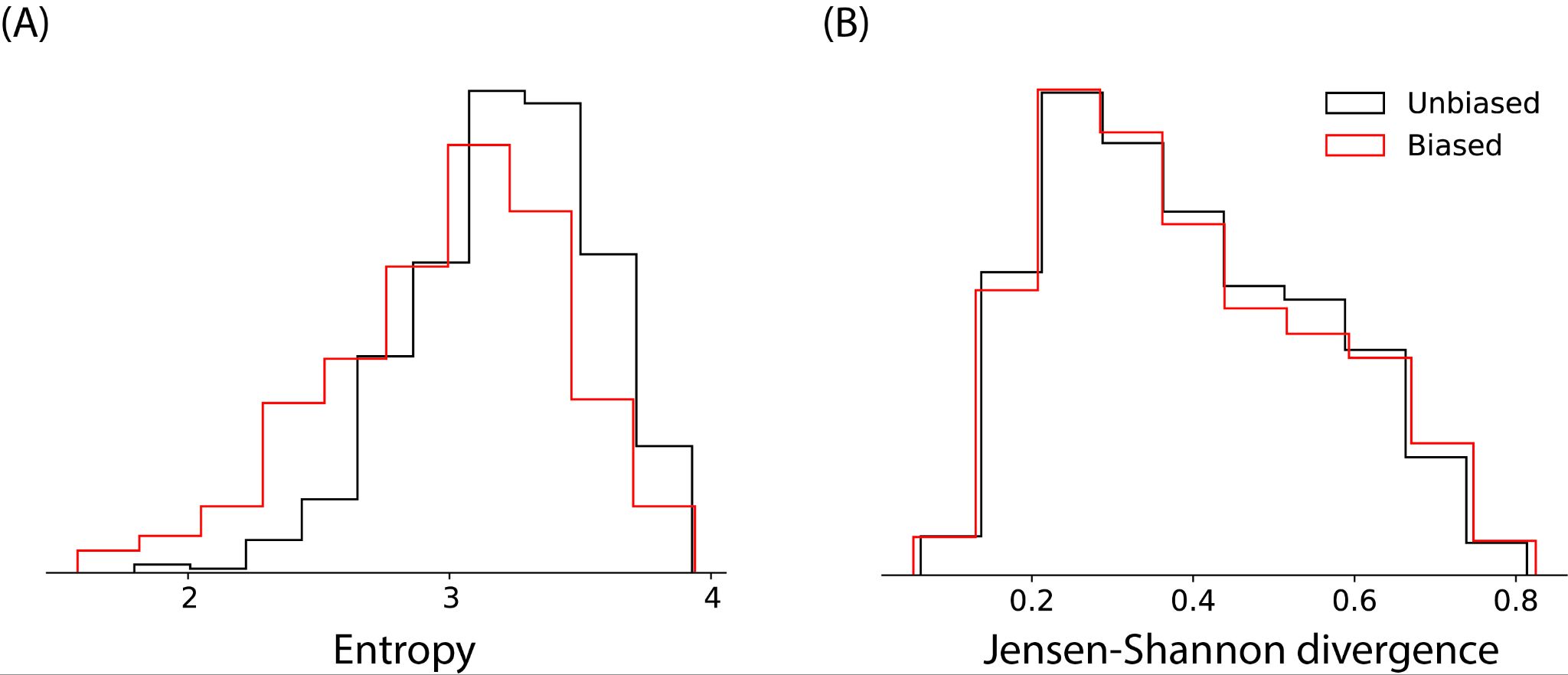

**SI figure 9.** A comparison between $\alpha-$ and $\beta-$diversity and species abundance distribution for naturally occurring microbiomes and microbiomes that correspond to high pH and high starch intake.
